# Photo-Disassembly of Membrane Microdomains Revives Conventional Antibiotics against MRSA

**DOI:** 10.1101/747626

**Authors:** Jie Hui, Pu-Ting Dong, Lijia Liang, Taraknath Mandal, Junjie Li, Erilinda R. Ulloa, Yuewei Zhan, Sebastian Jusuf, Cheng Zong, Mohamed N. Seleem, George Y. Liu, Qiang Cui, Ji-Xin Cheng

## Abstract

Confronted with the rapid evolution and dissemination of antibiotic resistance, there is an urgent need to develop alternative treatment strategies for drug-resistant pathogens. Here, we present an unconventional approach to restore the susceptibility of methicillin-resistant *S. aureus* (MRSA) to a broad spectrum of conventional antibiotics via photo-disassembly of functional membrane microdomains. The photo-disassembly of microdomains is based on effective photolysis of staphyloxanthin, the golden carotenoid pigment that gives its name. Upon pulsed laser treatment, cell membranes are found severely disorganized and malfunctioned to defense antibiotics, as unveiled by membrane permeabilization, membrane fluidification, and detachment of membrane protein, PBP2a. Consequently, our photolysis approach increases susceptibility and inhibits development of resistance to a broad spectrum of antibiotics including penicillins, quinolones, tetracyclines, aminoglycosides, lipopeptides, and oxazolidinones.

**One Sentence Summary:** Using photons to crash *S. aureus* cell membrane and its formidable defense against a broad spectrum of antibiotics.

---

Antibiotic resistance in human pathogens is one of the biggest public health challenges of our time. One such deadly pathogen is *S. aureus* or particularly methicillin-resistant *S. aureus* (MRSA), which causes high morbidity and mortality worldwide. An estimate of 23,000 fatalities occur each year in US due to antibiotic-resistant infections; surprisingly, nearly half of these deaths (11,285) is due to one bacterial pathogen, MRSA (*1*). The prevalence of its antibiotic resistance is consistently challenging our current treatment options via various molecular mechanisms. Particularly, overexpression of *mecA* encoded penicillin-binding protein 2a (PBP2a) in MRSA strains reduces the affinity of most beta-lactams (*2, 3*); active efflux pumps on cell membranes keep intracellular antibiotic concentration at sublethal level, conferring multi-drug resistance to fluoroquinolones and tetracyclines (*4, 5*); remodeling of membrane composition, e.g. phospholipids, reduces the binding thus the effectiveness of daptomycin, a last-resort antibiotic (*6*). Moreover, the development of new antibiotics is currently unable to keep pace with the emergence of resistant bacteria, thus likely leading us to a post-antibiotic era (*7*). To tackle this grand challenge, alternative treatment strategies are urgently required.

Grounded on the increasing understanding of virulence factors in disease progression and host defense, anti-virulence strategies have arisen in the past decade as an alternative (*8-10*). In *S. aureus*, staphyloxanthin (STX), the yellow carotenoid pigment that gives its name, is a key virulence factor (*11*). This pigment is expressed for *S. aureus* pathogenesis and used as an antioxidant to neutralize reactive oxygen species (ROS) produced by the host immune system (*12*). Recent studies on cell membrane organization further suggest that STX and its derivatives condense as the constituent lipids of functional membrane microdomains (FMM), endowing membrane integrity and providing a platform to facilitate protein-protein oligomerization and interaction, including PBP2a, to further promote cell virulence and antibiotic resistance (*13*). Therefore, blocking STX biosynthesis pathways has become an innovative therapeutic approach. Thus far, cholesterol-lowering drugs, including compound BPH-652 and statins, have shown capability of inhibiting *S. aureus* virulence by targeting the enzymatic activity, e.g. dehydrosqualene synthase (CrtM), along the pathway for STX biosynthesis (*13, 14*). However, these drugs suffer from off-target issues, as human and *S. aureus* share the same pathway for biosynthesis of presqualene diphosphate, an intermediate used to produce downstream cholesterol or STX. Additionally, anti-fungal drug, naftifine, was recently repurposed to block STX expression and sensitize *S. aureus* to immune clearance (*15*). Despite these advances, all of these are still drug-based approaches to inhibit STX virulence, which require additional treatment time, accompany with serious side effects, show weak activities, and have higher risk for resistance development by targeting a single upstream biosynthetic enzyme, which will eventually prevent their clinical utilization.

In this study, we unveil that staphyloxanthin is the molecular target of photons within the entire blue wavelength range, demonstrating an unconventional way to deplete STX photochemically. Grounded on the STX photolysis kinetics, a short-pulsed blue laser was further identified to strip off this pigment with high efficiency and speed in wide field. In contrast to drug-based approaches, this photonic approach depletes the final product, STX, swiftly in a drug-free manner. More significantly, this disruption, enabled by the pulsed laser, fundamentally disorganizes and further malfunctions FMM as unveiled by increased membrane fluidity, ample membrane permeability, and PBP2a protein detachment, simultaneously and immediately after exposure. These membrane damages inhibit PBP2a deactivation of penicillins and facilitate the intracellular delivery and membrane insertion of conventional antibiotics, specific to their mechanisms of action. As a result, photo-disassembly of FMM restores the susceptibility and inhibits resistance development to a broad spectrum of conventional antibiotics against MRSA. Additionally, this work further deciphers the structural and functional properties of STX-enriched membrane microdomains for antibiotic resistance, thus providing a strategy to tackle antibiotic resistance by targeting STX virulence.

## Results

### Pulsed blue laser photolysis of staphyloxanthin

In order to test the hypothesis that STX is the molecular target of photons in the entire blue range, we directly exposed high-concentration stationary-phase MRSA colony to a wavelength-tunable laser beam in a wide-field illumination configuration as shown in **Fig. 1A**. Strikingly, the distinctive golden color of MRSA colony fades quickly over time when the wavelengths were tuned into the blue (400-490 nm) wavelength range (e.g. the images of MRSA colony with 460 nm illumination wavelength in **Fig. 1A**). As the golden colony color is originated from STX pigment, such color-fading phenomenon suggests that STX is subject to photolysis (molecular structure of STX shown in **Fig. 1A**). In order to further validate this point, we applied resonance Raman spectroscopy to quantify STX content in MRSA cells by taking advantage of its high sensitivity, molecular specificity, and linear concentration dependence (*16*). STX in MRSA shows three characteristic Raman peaks around 1008 (methyl rocking), 1161 (C-C stretch), and 1525 cm^-1^ (C=C stretch), respectively, corresponding to their specific molecular vibrational modes (**Fig. 1B**). With increased laser treatment time, we observed dramatically decreased peak amplitude for all three Raman bands, suggesting the cleavage of both C-C and C=C bonds that constitutes the polyene chain of STX (**Fig. 1B**). As a result, the unsaturated tail of STX, the nine conjugated C=C double bonds, is decomposed or truncated, as confirmed by mass spectrometry (*17*). In contrast, when we blocked STX biosynthesis in *S. aureus* by knocking down CrtM, namely *S. aureus* ΔCrtM, its colony turns colorless and shows no detectable peaks for all three Raman bands, confirming that these Raman bands are exclusively from STX (**Fig. 1C**). With fixed laser power and dosage (50 mW, 5 min exposure time), MRSA colonies were further illuminated at different laser wavelengths and STX photolysis efficiency calculated using the Raman peak amplitude at 1161 cm^-1^ before and after illumination. The results in **Fig. 1D** indicate that STX is subject to effective photolysis in the entire blue wavelength range (400-490 nm) with significantly reduced efficiency when above 500 nm. This efficiency curve matches the absorption spectrum of STX as photolysis is grounded on the absorption of chromophores (**fig. S1A**). The effective STX photolysis induces significant absorption change, which is directly reflected on the absorption spectra of MRSA bacterial solution (**fig. S1A**). By compromising the STX photolysis efficiency and optical penetration, 460-480 nm is the preferable optical window (460 nm illumination wavelength was applied in the following studies). Notably, STX photolysis behavior is not only limited to MRSA, but broadly shown on vancomycin-resistant *S. aureus* (VRSA) and other clinically isolated multi-drug resistant *S. aureus* strains (**fig. S1B** and **Fig. 1E**; their minimum inhibitory concentrations shown in **Table S1**), as more than 90% of all *S. aureus* human clinical isolates generate this golden pigment (*18*). Collectively, these results suggest that STX is the molecular target of photons or lasers in the entire blue range.

**Fig. 1.**
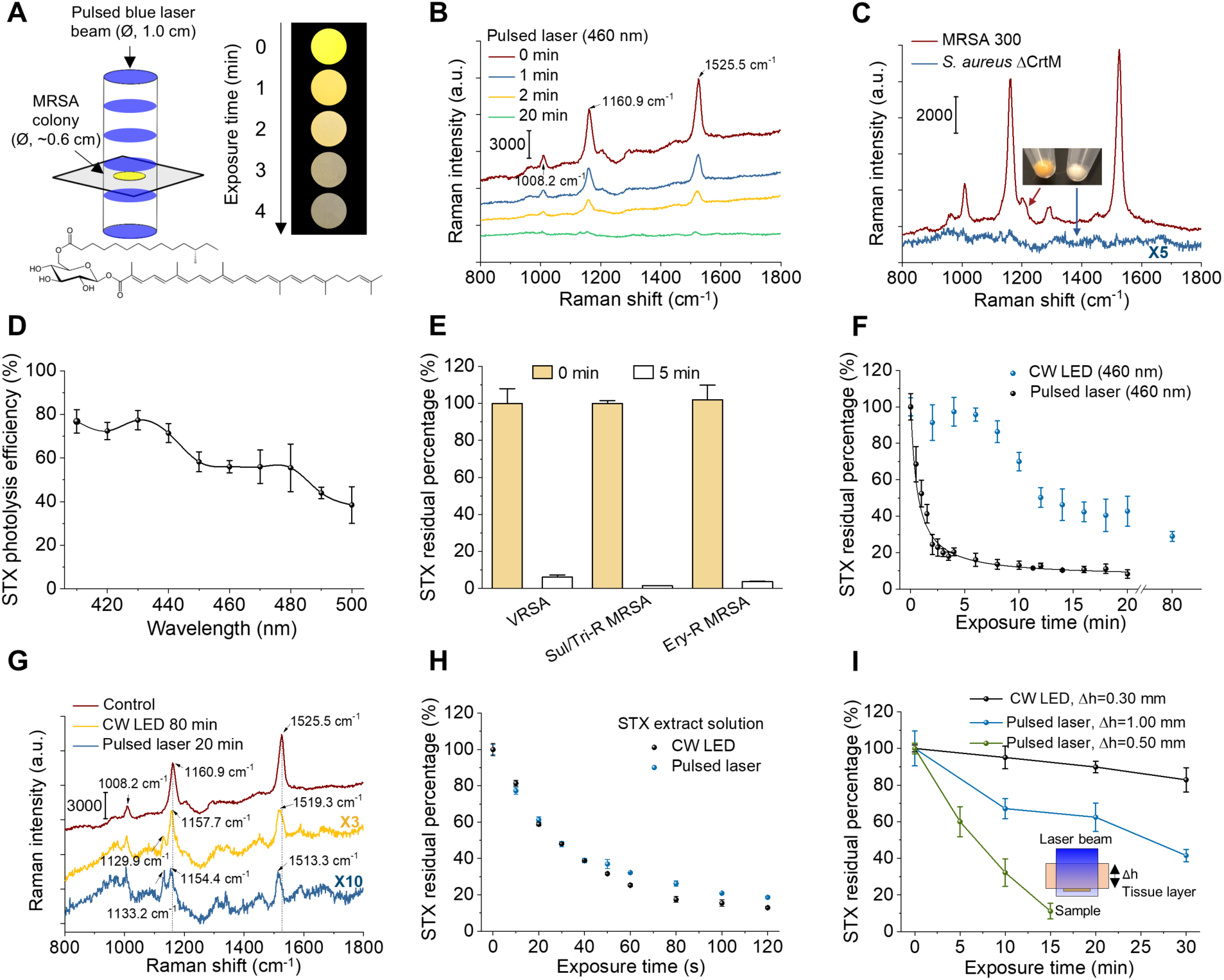
Photophysics and photochemistry of pulsed laser photolysis of STX. (A) (Left) Schematic of MRSA colony (or MRSA solution or STX extract solution) treated by nanosecond pulsed laser in a wide-field illumination configuration. (Right) Digital images of MRSA colony over laser treatment time to show golden color fading phenomenon. Image were recorded with sample placed on a transparent glass cover slide over a black paper. (Bottom) STX molecular structure. Ø refers diameter of bacterial colony. (B) Resonance Raman spectroscopy of MRSA colony over 460 nm nanosecond pulsed laser treatment time (measured on the same colony). Numbers indicate major Raman peak positions. (C) Resonance Raman spectroscopy of MRSA and *S. aureus* ΔCrtM colonies. The images show the color of spun-down cells. (D) Spectroscopic study of STX photolysis efficiency with nanosecond pulsed laser power of 50 mW and treatment time of 5 min. STX photolysis efficiency is quantified by Raman peak amplitude at 1161 cm^-1^. (E) Raman quantification of STX abundance in multidrug-resistant *S. aureus* cells before and after 5 min laser treatment (460 nm). Bacterial strains include vancomycin-resistance *S. aureus* (VRSA), sulfamethoxazole/trimethoprim-resistant MRSA (Sul/Tri-R MRSA), and erythromycin-resistant MRSA (Ery-R MRSA). (F) STX photolysis kinetics of MRSA colony by nanosecond pulsed laser and CW LED under the same illumination power, area, and center wavelength (460 nm). Solid black curve is the fitting result by a second-order photobleaching model. (G) Resonance Raman spectroscopy of STX in MRSA colony with or without long time-treatment by nanosecond pulsed laser and CW LED at 460 nm highlighting STX photolysis induced Raman peak shifts and the generation of new Raman peak. Numbers indicate Raman peak positions before and after light treatment. (H) STX photolysis kinetics of STX solution by nanosecond pulsed laser and CW LED under the same illumination power, area, and center wavelength (460 nm). STX solution were extracted directly from MRSA cells. (I) STX photolysis kinetics of MRSA colony placed beneath a tissue layer with various thickness by nanosecond pulsed laser and CW LED under the same illumination power, area, and center wavelength (460 nm). The inset shows the schematic of experimental scheme. Δh indicates the thickness of tissue layer. CW, continuous wave. The cells used were all cultured to reach 3-day stationary phase. N=3 for all the above measurements.

Considering the significance of STX virulence in a MRSA-caused disease, an optimal light source that enables efficient, fast, complete, and deep depletion of STX is of great importance. Our previous study via transient absorption microscopy suggests that STX photolysis under tightly focused laser primarily follows a second-order photolysis model due to triplet-triplet annihilation: T* + T* → R + S, where R and S represent reduced and semi-oxidized forms (*17*). The triplet excitons form with high yield via singlet fission when carotenoids self-assemble into multimer or aggregates on cell membrane (*19*). As the triplet lifetime of STX is on a microsecond scale (*20*) and STX laterally assembles within FMM (*13*), a high-fluence nanosecond pulsed laser can be used to effectively populate STX molecules to their triplet state within single pulse excitation thus accelerating STX photolysis nonlinearly.

To test this hypothesis, we firstly exposed stationary-phase MRSA colony to the nanosecond pulsed laser and a continuous-wave light-emitting diode (LED) with output power of 120 mW, with wavelength centered around 460 nm, then monitored their residual STX through resonance Raman spectroscopy over different exposure time. Remarkably, the nanosecond pulsed laser shows unmatched efficiency, speed, and completeness for STX photolysis when compared with the LED, as it depletes 80% of STX in MRSA cells within less than 2 mins, whereas it takes LED more than 20 mins to reach the same efficiency (**Fig. 1B,F** and **fig. S1C**); the STX photolysis by LED is not complete even over 80 mins illumination. The efficiency and speed come from the nonlinearity of STX annihilation enabled by nanosecond pulsed laser, consistent with the second-order fitting (*17*) result of the decay curve (**Fig. 1F**). By closely examining the Raman spectra, nanosecond pulsed laser further induces significant blue shifts of these peaks; the shifts are as large as 12 and 6 cm^-1^ for peaks at 1525 and 1161 cm^-1^, respectively (**Fig. 1G**). These blue shifts provide additional evidence to support the photochemistry process in STX. In contrast, when we monitored the photolysis kinetics on STX solution extracted from MRSA pellets, nanosecond pulsed laser and LED no longer show distinctive decay curves (**Fig. 1H** and **fig. S1D,E**). Therefore, STX photolysis speed as suggested has high concentration dependence; highly aggregated STX nonlinearly increases STX photolysis efficiency and speed. When laser pulse fluence was doubled meanwhile keeping illumination dosage the same, photolysis delay curves for nanosecond pulsed laser only show minor difference, as likely this 2-time difference in pulse fluence is minor when compared with 10^7^-time difference between nanosecond pulsed laser and continuous-wave LED under the same power (**fig. S1F**). Thus, further shortened illumination time can be achieved by simply increasing pulse fluence until reaching saturation. More significantly, the high-fluence nanosecond pulsed laser enables ∼4-fold larger treatment depth when compared with LED, as more than 50% STX molecules are depleted by nanosecond pulsed laser when MRSA colonies are placed beneath a tissue layer with thickness beyond 1 mm within one cell cycle (30 mins), whereas LED barely penetrates through 300 µm tissue to reach the same efficiency (**Fig. 1I**, experimental schematic shown in the inset). Such effective STX photolysis in deep tissue comes from the conjugation of the photolysis nonlinearity and high photon fluence of pulsed laser, as the photons fluence of pulsed laser is several orders of magnitude higher than that of low-level light sources (e.g. LED) even through a thick tissue layer. The extended depth is sufficient to penetrate and treat MRSA biofilms (thickness typically ranging from a few micrometers to several hundreds of micrometers (*21*)), which are normally difficult to treat by antibiotics due to biofilm-mediated inactivation (*22*). Notably, the power and dosage for nanosecond pulsed blue laser in this study are below the American National Standards Institute (ANSI) safety limit for human skin exposure to lasers at 460 nm (*23*). In contrast to continuous-wave LED, nanosecond pulsed laser further eliminates potential photothermal issues as the temperature rise on human skin is quite small (<5 degree). Collectively, these results suggest that high-fluence short-pulsed blue laser is the superior light source to deplete STX in MRSA quickly, effectively, completely, and safely.

### Photo-disassembly of FMM: membrane permeabilization

STX is known acting as the constituent lipid of FMM, which are embedded in the lipid bilayer of virulent *S. aureus* strains and implicated in maintenance of membrane integrity. Therefore, we hypothesize that STX photolysis disrupts membrane integrity by increasing membrane permeability, thus facilitating the intracellular accumulation of small-molecule dyes or antibiotics via passive diffusion (**Fig. 2A**). To prove this point, membrane permeability with or without laser treatment was evaluated in real time by SYTOX green (600 Da), a fluorescent dye for nucleic acids stain of cells only with compromised membrane. With increased laser treatment time, a significantly larger and faster uptake of SYTOX green is observed, indicating severely compromised cell membranes; whereas cells without laser treatment show negligible uptake, which validates the role of STX on membrane integrity (**Fig. 2B**). These results were further confirmed by confocal fluorescence imaging and statistical analysis of signal intensity for individual cells. From **Fig. 2C,D** and **fig. S2A**, significantly brighter fluorescence signal from the entire cell population is observed over laser treatment time, indicating different levels of membrane permeability. After 10 min laser treatment, such damaged membranes are unable to recover even with 2 hours culturing (**fig. S2B**). In contrast, for *S. aureus* with ΔCrtM (nonpigmented mutant) and log-phase MRSA, no significant difference in SYTOX green uptake is shown between laser treated vs the untreated (**Fig. 2E** and **fig. S2C**).

**Fig. 2.**
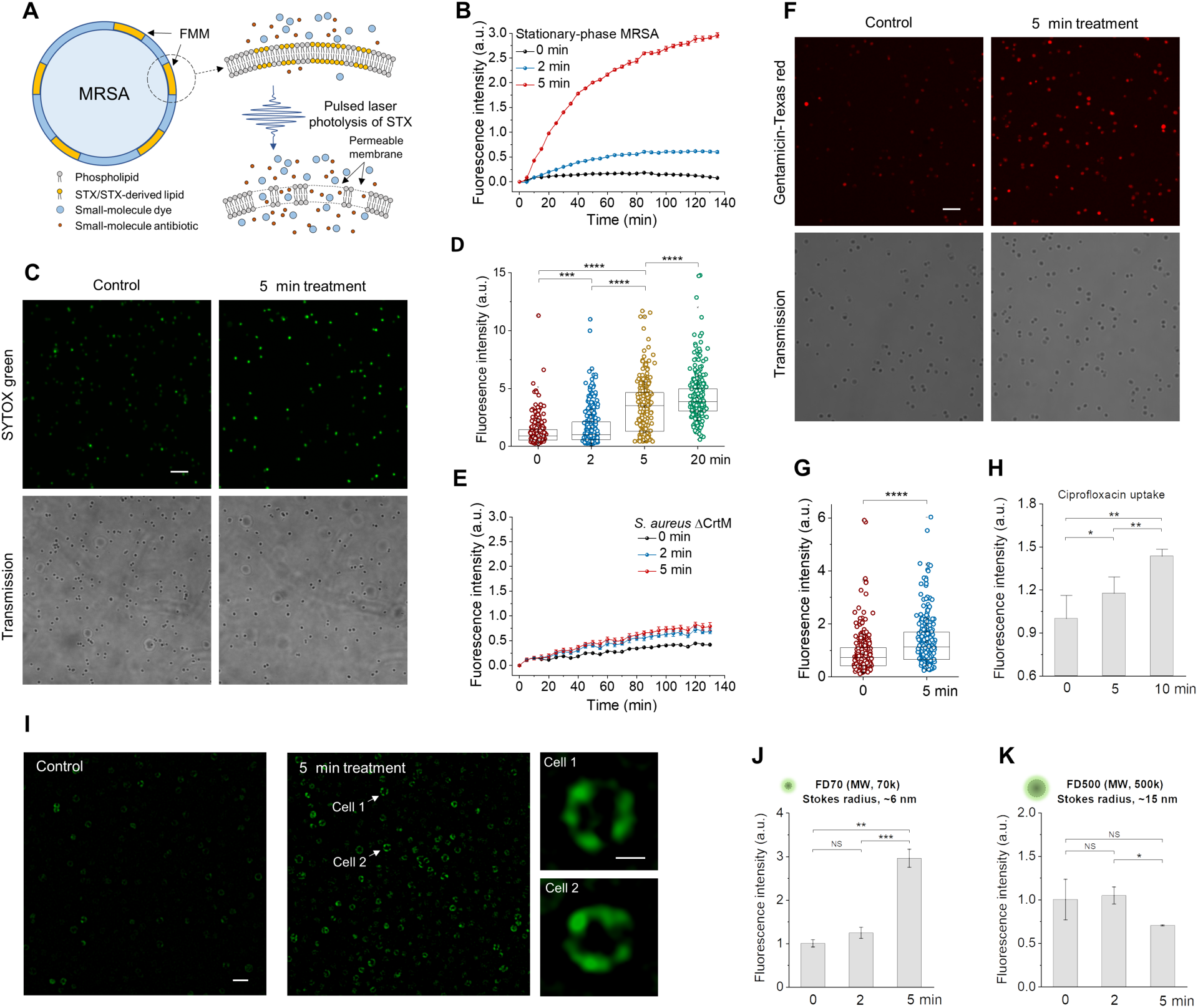
First mechanism for photo-disassembly of membrane microdomains: membrane permeabilization. (A) Schematic of membrane permeability mechanism via pulsed laser photolysis of STX. (B) Real-time intracellular uptake kinetics of SYTOX green by stationary-phase MRSA with or without pulsed laser treatment. (C) Confocal fluorescence images of intracellular uptake of SYTOX green by stationary-phase MRSA cells with or without pulsed laser treatment. (Top) fluorescence images. (Bottom) corresponding transmission images. (D) Statistical analysis of fluorescence signal from MRSA cells in (C) from each treated group with N ≥ 300 per group. (E) Real-time intracellular uptake kinetics of SYTOX green by stationary-phase *S. aureus* ΔCrtM with or without pulsed laser treatment. (F) Confocal fluorescence images of intracellular uptake of gentamicin-Texas red by stationary-phase MRSA cells with or without pulsed laser treatment. (Top) fluorescence images. (Bottom) Corresponding transmission images. (G) Statistical analysis of fluorescence signal of MRSA cells in (F) from each treated group with N ≥ 300 per group. (H) Fluorescence detection of ciprofloxacin uptake by stationary-phase MRSA with or without pulsed laser treatment. (I) Structured illumination microscopic images of FD500 uptake by stationary-phase MRSA cells with or without pulsed laser treatment. Insets shows representative images of FD500 distribution on single cell after 5 min laser treatment. Fluorescence detection of (J) FD70 and (K) FD500 uptake by stationary-phase MRSA with or without pulsed laser treatment. MW, molecular weight. Scale bar, 5 µm for (C, F, I) and 0.5 µm for zoom-in images in (I). N=3 for all the above measurements.

Based on these findings, we further hypothesize that increased membrane permeability induced by STX photolysis would allow passive diffusion of small-molecule antibiotics that target intracellular activities. To demonstrate this point, we used the aminoglycoside, gentamicin, as an example. Gentamicin was firstly conjugated with a fluorescent dye, Texas red, and then imaged via confocal fluorescence microscopy after co-culturing with cells. As expected, cells with laser treatment accumulate significantly more gentamicin molecules than untreated, from either single cells (**Fig. 2F,G**) or the entire cell population (**fig. S2D**). The uptake of ciprofloxacin, another small-molecule antibiotic that belongs to fluoroquinolone class, can be directly detected via its endogenous fluorescent nature. Compared to the untreated cells, increased fluorescence signal is shown on cells with laser treatment (**Fig. 2H**). These results further confirm that small-molecule antibiotics can diffuse into the cell via permeable membrane induced by laser treatment.

To estimate how large a molecule can diffuse into the damaged membrane, we applied dextran labeled fluorescein isothiocyanate (FITC-dextran) with variable molecular weight/Stokes radius and monitored its insertion before and after laser treatment. For FD70 with molecular weight of 70k Da and Stokes radius of 6 nm, longer laser treatment time yields increased fluorescence signal either at individual cell level (**Fig. 2I**) or from total cell population (**Fig. 2J**). Laser treatment over 5 min leads to ample insertion of FD 70 (**Fig. 2J**). Super-resolution imaging of individual cells further shows that these dyes are primarily inserted and concentrated within FMM (**Fig. 2I**, zoom-in images). In contrast, when FD500 with molecular weight of 500k Da and Stokes radius of 15 nm was applied, no uptake is shown, indicating an upper limit on molecular Stoke radius of 30 nm level (**Fig. 2K**). These results suggest that after effective STX photolysis, FMM becomes porous, allowing molecules with Stokes radius up to nanometer level to diffuse through or insert into the membrane.

### Photo-disassembly of FMM: membrane fluidification

After effective STX photolysis, its products no longer maintain the chemical structure and properties of STX. The unsaturated tail of STX is truncated as unveiled by Raman spectroscopy results; the polarity of its products becomes significantly higher than that of STX as suggested by liquid chromatography results (*17*). As a result, these products spontaneously tend to disperse or detach from their original membrane organization. These behaviors profoundly disrupt the lipid packing within the microdomain, thus changing the membrane fluidity and subsequently facilitating the insertion of membrane targeting antibiotics, e.g. daptomycin. To test this hypothesis, we evaluated the membrane fluidity with or without laser treatment by DiIC_18_, a fluorescent dye that displays affinity for membrane areas with increased fluidity due to its short hydrocarbon tail (*24*) (**Fig. 3A**). As shown in **Fig. 3B,C**, significantly more DiIC_18_ is shown up as foci in log-phase MRSA when compared to the stationary-phase, as membrane in stationary phase becomes more rigid than that in log phase, partially due to the presence of rigid STX (*25*). After laser treatment, the foci number on each cell is significantly increased when compared with that of stationary-phase cells without laser treatment. Notably, 70% of cells in stationary phase show no detectable fluorescence signal, whereas this portion drops dramatically to 35% after 2.5 min laser treatment. The ample uptake indicates that laser treatment renders membrane more fluid due to the depletion of rigid unsaturated STX tail and the subsequent loose packing of lipid bilayer.

**Fig. 3.**
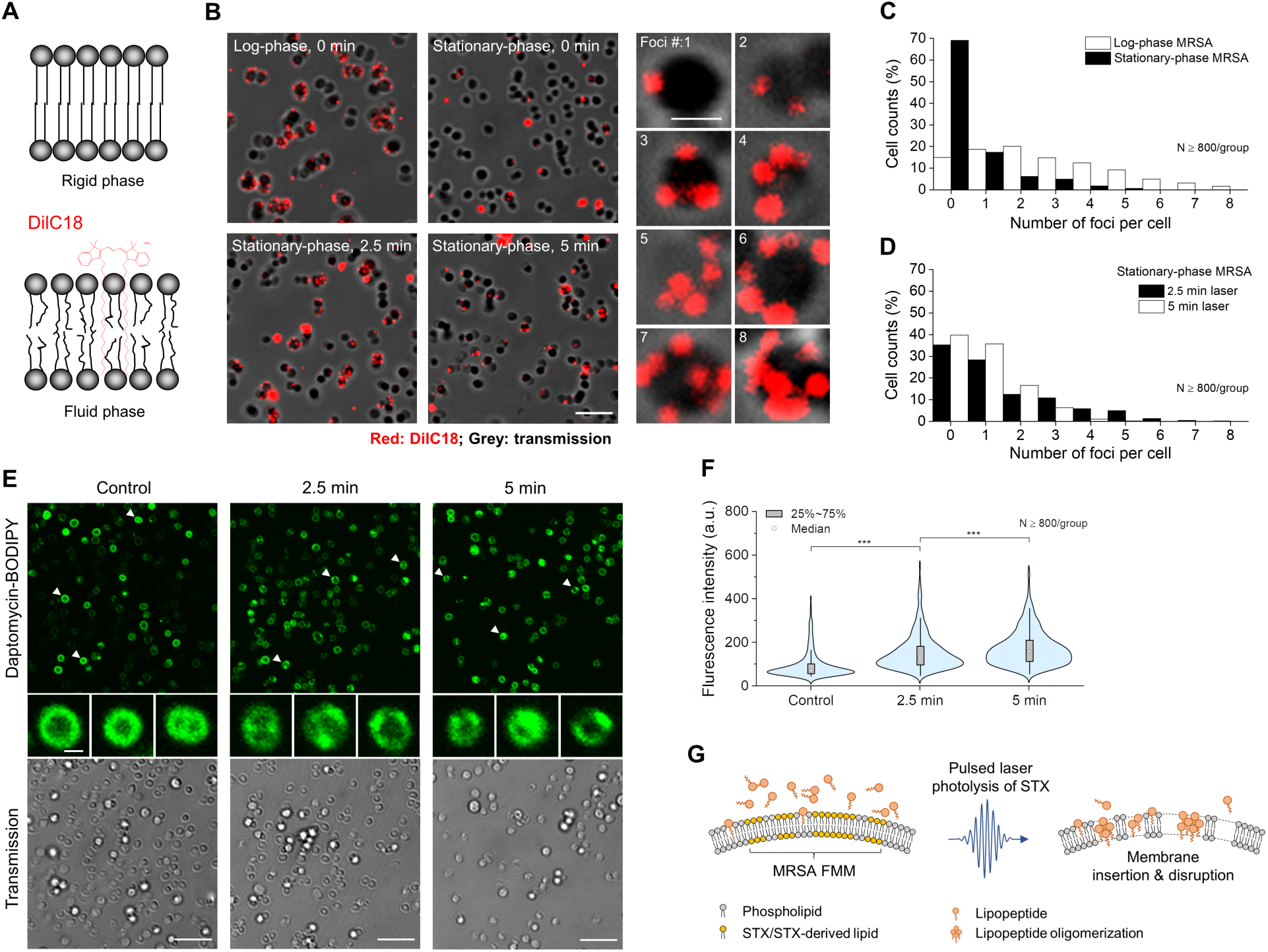
Second mechanism for photo-disassembly of membrane microdomains: membrane fluidification. (A) Schematic of membrane insertion of DiIC_18_ induced by gel/rigid-to-liquid/fluid phase change. (B) (Left and middle columns) fluorescence images of DiIC_18_ foci formation for groups including log-phase MRSA and stationary-phase MRSA with or without laser treatment. (Right column) zoom-in fluorescence images of MRSA cells with different foci number on each cell. Fluorescence from DiIC_18_, red; transmission, grey. Scale bar, 5 µm for (left and middle columns) and 1 µm for (right column). Statistical analysis of foci number on cells from each group in (B): (C) log-phase and stationary-phase MRSA without laser treatment; (D) stationary-phase MRSA with different laser treatment time. N ≥ 800/group. (E) (Top row) fluorescence images of daptomycin-BODIPY on stationary-phase MRSA with or without laser treatment. (Middle row) representative zoom-in images of the upper row. (Bottom row) corresponding transmission channels. Scale bar, 5 µm for (top and bottom row) and 0.5 µm for (middle row). (F) Statistical analysis of fluorescence signal intensity from MRSA cells in (E) with or without laser treatment with N ≥ 800. (G) Schematic of antibiotic membrane insertion mechanism via pulsed laser photolysis of STX.

The increased membrane rigidity by STX overexpression promotes the bacterial resistance against daptomycin, a cationic antimicrobial peptide, by reducing its membrane binding and subsequent membrane disruption (*25-27*). Therefore, we further hypothesize that increased membrane fluidity after STX photolysis facilitates the insertion of daptomycin. To prove this point, we first labeled daptomycin with BODIPY (molecular structure shown in **fig. S3A**), then imaged cellular uptake of daptomycin with or without laser treatment. From **Fig. 3E,F**, significantly more daptomycin uptake is shown for laser-treated groups when compared to the untreated groups; longer treatment yields higher uptake. More interestingly, daptomycin distribution between laser treated and untreated groups are quite different; for the untreated, daptomycin distributes evenly on the cell membrane, whereas, aggregates or domain-like structures with bright signal are found on cells after laser treatment (representative zoom-in images in the middle row in **Fig. 3E**). These aggregates most likely form within FMM due to the promoted insertion and oligomerization of daptomycin. Collectively, these results provide evidences to support the ample increase of membrane fluidity after STX photolysis, thus potentiate antibiotic lipopeptides to insert and oligomerize within the domains and further disrupt cell membrane as illustrated in **Fig. 3G**.

### Photo-disassembly of FMM: membrane protein detachment

To demonstrate how STX photolysis further malfunctions membrane proteins that are co-localized within STX-enriched FMM, we chose penicillin-binding protein 2a, PBP2a, as an example. MRSA acquires resistance to beta-lactam antibiotics through expression of PBP2a, a protein (*2*) that primarily anchors within FMM through its transmembrane helix and hides its targeting site inaccessible by beta-lactam antibiotics (**Fig. 4A**). Considering the relative structural organization of STX and PBP2a, we hypothesize that PBP2a protein complex can be disassembled and unanchored from cell membrane upon effective STX photolysis. To validate this point, we first resolved the structural distribution of PBP2a under a structured illumination microscopy via immunostaining with anti-PBP2a antibodies both for laser-treated (**Fig. 4B,C**) and the untreated (**Fig. 4D,E**). For the untreated, we observed bright fluorescence signal from all stationary-phase MRSA cells due to ample PBP2a expression. These proteins are accumulated discretely within small membrane domains as visualized in both 3-D (**Fig. 4B**) and 2-D along various depths (**Fig. 4C**). Three to four foci on average is found on each cell, indicating the prevalence of microdomain formation when cells reach their stationary phase (**fig. S4A**). Once treated with pulsed laser, dramatically decreased signal intensity and altered signal distribution are observed on each individual cell (**Fig. 4D,E**). Laser-treated cells have around 2 times lower signal intensity when compared with the untreated, thus indicating a large portion of PBP2a proteins are detached from cell membrane (**Fig. 4F**). The left PBP2a proteins are dispersed laterally with its dispersion quantified by coefficient of variation, which is significantly higher than that of the untreated (**Fig. 4G**, quantification method shown in **fig. S4B**). Such detachment and dispersion lead to significantly reduced contrast between FMM and its neighboring lipid bilayer (**Fig. 4E**). Western blotting results further confirms the PBP2a detachment mechanism, as increased amount of PBP2a is found in supernatant over laser treatment time, whereas decreased amount found in MRSA pellets (**Fig. 4H**). Taken together, photolysis of the constituent lipids leads to disassembly and detachment of PBP2a from FMM, thus disables MRSA’s defense to penicillins as illustrated in **Fig. 4I**. Additionally, as PBP2a is primarily utilized to catalyze cell-wall crosslinking, their detachment further affects cell wall synthesis and potentially cell viability.

**Fig. 4.**
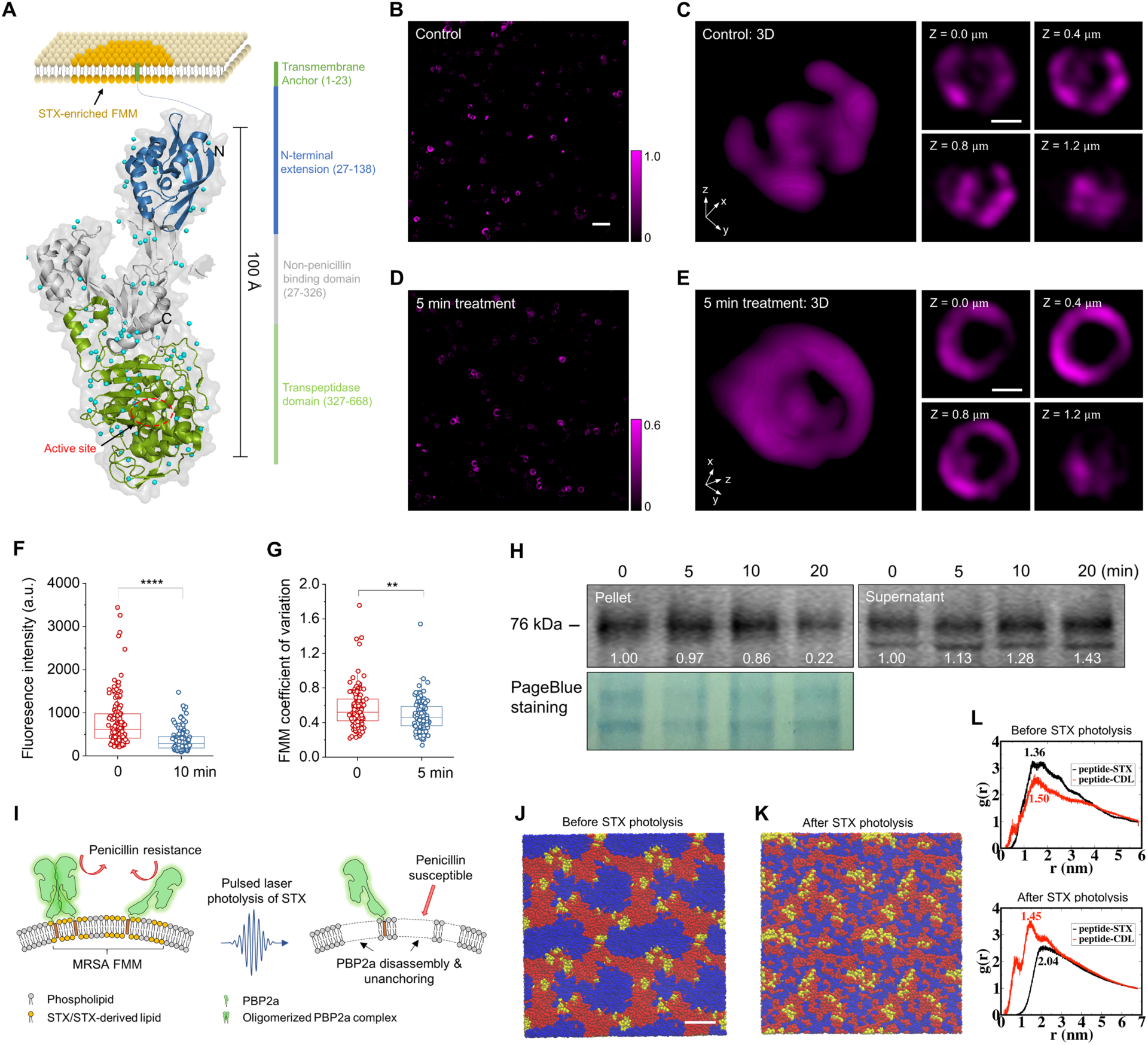
Third mechanism for photo-disassembly of membrane microdomains: membrane protein detachment. (A) Schematic of PBP2a protein structure and location relative to STX enriched membrane microdomain. (B, C) SIM images of PBP2a via immunostaining on MRSA cells in (B) 2-D and (C) 3-D. Intensity color bar applies to (B, C). (D, E) SIM images of PBP2a immunostaining on MRSA cells in (D) 2-D and (E) 3-D after 5 min laser treatment. Intensity color bar applies to (D, E). Scale bar, 2.0 µm for (B, D) and 0.5 µm for (C, E). (F) Statistical analysis of signal intensity from MRSA cells with or without laser treatment with N ≥ 100. (G) Statistical analysis of PBP2 coefficient of variation on MRSA cells with or without laser treatment with N ≥ 100. (H) Western blot of PBP2a on MRSA pellets and its supernatant for groups with different laser treatment time. Numbers indicate the integrated signal intensity. Pageblue staining of the same samples was used as a loading control. (I) Schematic of PBP2a disassembly and detachment mechanism via pulsed laser photolysis of STX. (J) Self-assembled microphase separated domain structures of modeled membrane after 10 µs molecular dynamics simulation. Full-length STX lipids, red; cardiolipin lipids, blue; PBP2a peptides, yellow. (K) Final configuration of modeled membrane with truncated STX after 10µs molecular dynamics simulation. Color scheme also applies to (J). Water and ions are made invisible for clarity for (J, K). Scale bar, 5 nm for (J, K). (L) RDFs of PBP2a peptides relative to the full-length STX and cardiolipins (upper panel) and truncated STX and cardiolipins (lower panel). Numbers on the plot indicate the locations of the first peak for each RDF.

To further investigate the membrane phase and its mechanical properties, we built a coarse-grained membrane model that contains STX, cardiolipin lipids, and transmembrane helixes of PBP2a proteins (coarse-grained representations shown in **fig. S4C-F**) and performed microsecond-scale molecular dynamics simulations. At the initial simulation configuration, STX, cardiolipin lipids and peptides randomly disperse in the built bilayer (**fig. S4G**). During 10 µs simulation, these molecules spontaneously self-assemble to a microphase separated system containing well distinguishable STX and cardiolipin microdomains, despite cardiolipin being a charged lipid; PBP2a peptides localize to the center of STX domains or the vicinity of STX/cardiolipin domain interface (**Fig. 4J**). The formation of microdomain is primarily driven by the preferable interactions among lipid tails of similar saturation or unsaturation nature, as in current system all four tails of cardiolipin are saturated, whereas STX lipid has a long unsaturated tail. This result is consistent with lipid domain formation commonly found for systems with a mixture of saturated and unsaturated lipids such as DOPC/DPPC, DOPC/DPPG, DOPG/DPPC and many others (*28*). To quantify the relative position and abundance of PBP2a peptides relative to STX and cardiolipin lipids, the radial distribution functions (RDFs), *g*(*r*), of PBP2a peptides were calculated. **Figure 4L** (upper panel) shows that the RDF peak of PBP2a peptide to STX is higher and located at smaller distance when compared to that of PBP2a peptide to cardiolipin, indicating that PBP2a peptides preferentially interact with STX lipids over cardiolipin, due likely to the better packing between the rigid fully unsaturated STX tail and the PBP2a transmembrane helix.

Our Raman spectroscopy results suggest that photolysis of STX leads to the loss of its rigid and unsaturated tail, the conjugated C=C chain. Thus, to mimic the scenario after STX photolysis, we repeated our simulations by replacing full-length STX with truncated STX with its unsaturated tail removed from the model (**fig. S4D**). Interestingly, the truncated STX lipids no longer form microdomains. As a result, all the lipids and PBP2a peptides are randomly dispersed (**Fig. 4K**). Moreover, the RDF of PBP2a peptide to cardiolipin now features a higher peak at a smaller distance than that of PBP2a peptide to STX, suggesting that the PBP2a proteins prefer to interact with cardiolipins over truncated STX (**Fig. 4L**, lower panel). The different phase features before and after STX photolysis also lead to different membrane mechanics. For example, the calculated area expansion modulus (*K*_*A*_) of the membrane after microdomain formation is ∼58 *k*_*B*_*T*/*nm*^2^, which is significantly higher than the value of ∼42 *k*_*B*_*T*/*nm*^2^ with truncated STX, cardiolipins and peptides randomly dispersed after STX photolysis. This suggests that following the truncation of the unsaturated STX tail, the membrane loses the microphase separated domain structure and becomes more loosely packed, which in turn likely reduces the affinity of PBP2a protein to the membrane. Collectively, our simulations provide a plausible rational for the STX photolysis induced membrane remodeling, including the loss of functional domains, the increase of membrane permeability and fluidity, and the detachment of PBP2a from the membrane.

### Restoration of conventional antibiotics

With cell membrane catastrophically damaged via STX photolysis, we further reasoned that both cell growth and cell viability are severely compromised by laser treatment alone. To test this point, time-killing assay in phosphate-buffered saline was firstly performed on stationary-phase cells with or without laser treatment. Compared with the untreated, laser-treated cells are killed quickly and efficiently due to their disassembled FMM and incapacity for recovery (**Fig. 5A**). The killing efficiency shows strong illumination dosage/time dependence; 16 min laser treatment yields nearly 5-log killing when compared to the untreated. In contrast, *S. aureus* ΔCrtM shows relatively negligible killing by laser treatment (**Fig. 5B**). These results confirm that STX photolysis induced membrane disruption is the underlying eradication mechanism. Additionally, its recovery ability after laser treatment was assessed via a post-exposure effect assay, similar to post-antibiotic effect (*29*), as an important way to establish the optimal dosing regimen. The post-exposure effect of stationary-phase MRSA, depending on STX expression condition and laser treatment time, reaches up to 6-9 hours, due primarily to the membrane disruption mechanisms (**Fig. 5C** and **fig. S5A**), whereas no significant post-exposure effect observed for log-phase MRSA (**fig. S5B**) or *S. aureus* ΔCrtM (**Fig. 5D**). This post-exposure effect indicates a very slow recovery for stationary-phase cells after laser treatment thus fewer doses required for patients, which is superior than the post-antibiotic effect of most antibiotics including oxacillin, ofloxacin, and gentamicin (< 1 hour post-antibiotic effect for all three antibiotics, **fig. S5C**). More significantly, STX photolysis-induced FMM disassembly can pave a new approach to sensitize these bacteria to conventional antibiotics, even by antibiotics presumed to have no activity against MRSA, such as penicillins.

**Fig. 5.**
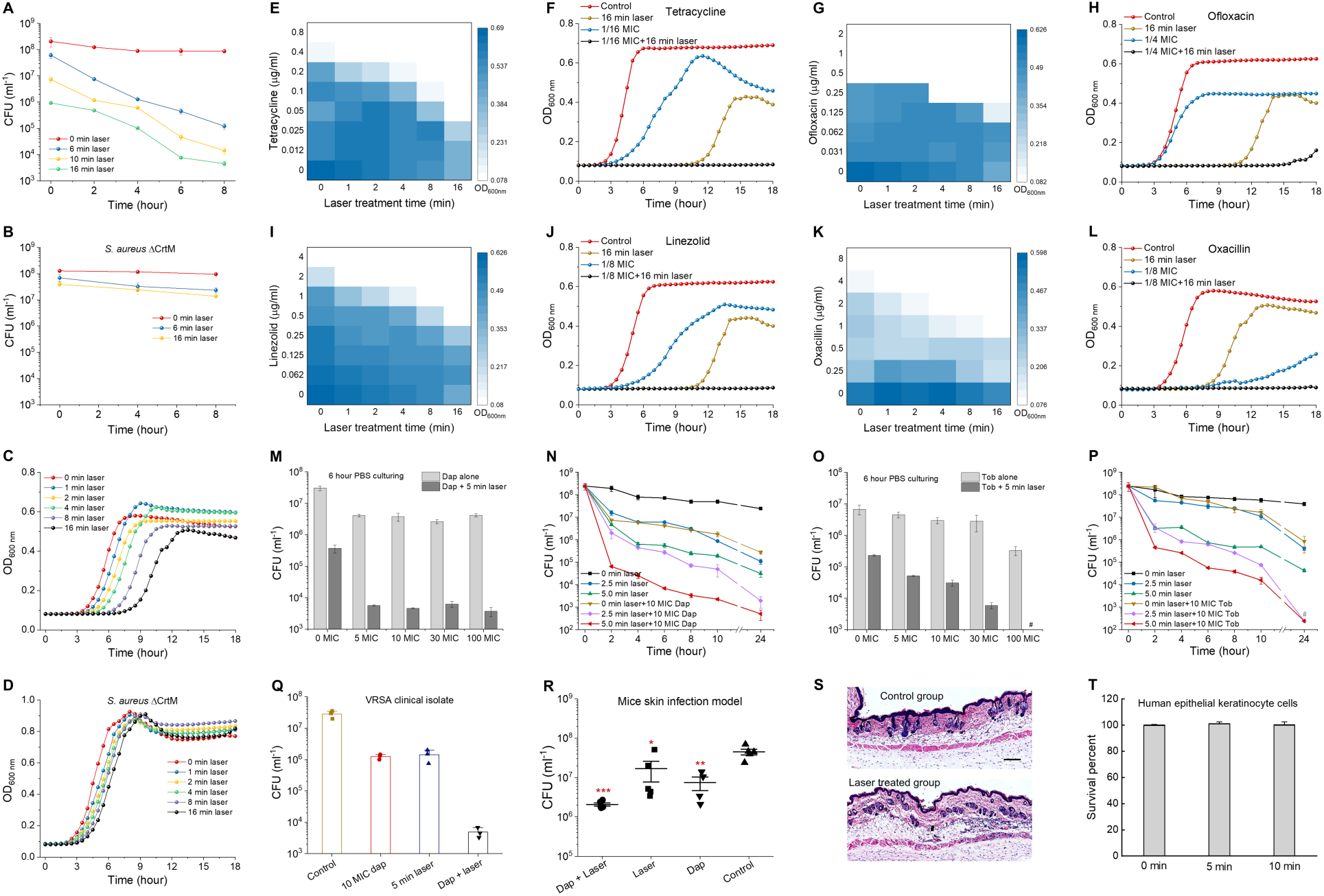
Photo-disassembly of membrane microdomains restores a broad spectrum of conventional antibiotics. Time-dependent killing of stationary-phase (A) MRSA and (B) *S. aureus* ΔCrtM cells in phosphate-buffered saline after different laser treatment time. Post-exposure effect of stationary-phase (C) MRSA and (D) *S. aureus* ΔCrtM after different laser treatment time. (E-L) Checkerboard assay results for synergy evaluation between laser treatment and different classes of antibiotics: (E, F) tetracycline, (G, H) ofloxacin, (I, J) linezolid, and (K, L) oxacillin. (F, H, J, L) Selected cell growth curves acquired from corresponding checkerboard assay results of each antibiotic. (M) Viability of stationary-phase MRSA after laser treatment alone or in combination with daptomycin with different concentrations followed by 6-hour incubation in phosphate-buffered saline. (N) Time-dependent killing of stationary-phase MRSA in phosphate-buffered saline after laser treatment alone or in combination with 10 MIC daptomycin. (O) Viability of stationary-phase MRSA after laser treatment alone or in combination with gentamicin with different concentrations followed by 6-hour incubation in phosphate-buffered saline. (P) Time-dependent killing of stationary-phase MRSA in phosphate-buffered saline after laser treatment alone or in combination with 10 MIC gentamicin. (Q) Time-dependent killing of stationary-phase vancomycin-resistant *S. aureus* (VRSA) strain in phosphate-buffered saline for four different treatment groups. (R) Efficiency of laser treatment alone or in combination with daptomycin on MRSA-caused mice skin infection model. (S) Hematoxylin and eosin stained histology evaluation of phototoxicity on mice skin. The mice used and treatment procedure applied were the same as that of (R) but without MRSA infection on the skin. (T) Viability of human keratinocyte cells over different laser treatment time to evaluate phototoxicity. N=5 for CFU enumeration for *in vivo* mice study. Dap, daptomycin; Tob, tobramycin. N=3 for the rest CFU enumeration, for checkerboard assay of each antibiotic and for phototoxicity evaluation on both human cells and *in vivo* mice.

To demonstrate the laser treatment-mediated synergism with antibiotics, we first applied the checkerboard assay as a screening method. Interestingly, synergism is identified between laser treatment and several major classes of antibiotics for MRSA growth inhibition (**Fig. 5E-L**). Using tetracycline as an example, the lowest concentration needed to completely inhibit MRSA growth within 18 hours is steadily decreased by elongated laser treatment time; 16 min laser treatment enables a 16-fold reduction, where 2-fold change or larger is regarded as synergy based on fractional inhibitory concentration index (FICI) (**Fig. 5E,F**). The similar results are found for quinolones: ofloxacin and ciprofloxacin (**Fig. 5G,H** and **fig. S5D,E**) and oxazolidinone: linezolid (**Fig. 5I,J**) with 2-fold, 8-fold reduction respectively. Notably, tetracyclines, oxazolidinones and quinolones all target intracellular activities; therefore, they have to penetrate through the membrane barrier in order to be functional. These growth inhibition results further validate our hypothesis that photo-disassembly of FMM renders membrane permeable to allow passive diffusion of small-molecule antibiotics inside cells, thus increasing their effectiveness against MRSA. Due to the disassembly and detachment of PBP2a proteins on cell membrane, laser treatment further re-sensitizes MRSA to penicillin: oxacillin with its concentration as low as 1 µg/ml, 8-fold lower than that of oxacillin-treated alone (**Fig. 5K,L**). In contrast, when vancomycin, an antibiotic that inhibits cell wall biosynthesis, was tested, no synergism is shown (**fig. S5F,G**). For bactericidal antibiotics, time-killing assay was then applied as the screening method. Due to laser-mediated membrane insertion and further disruption, 10-minimum inhibitory concentration (MIC) daptomycin is found capable of eradicating stationary-phase/dormant MRSA cells synergistically with only 5 min laser treatment (e.g. more than 3.5-log reduction after 6 hours), whereas antibiotics alone show very limited killing even at 100 MIC (e.g. 1-log reduction after 6 hours) (**Fig. 5M,N**). The similar synergistic killing is observed for aminoglycoside: tobramycin (**Fig. 5O,P**) due to its passive diffusion via laser-mediated permeable membrane. The synergistic therapy between 10 MIC daptomycin and laser treatment are also effective in eradicating VRSA and multidrug-resistant MRSA clinical isolates (**Fig. 5Q** and **fig. S5H,I**). Additionally, the synergy with laser treatment for MRSA killing is not only limited to conventional antibiotics; laser treatment facilitates human whole blood by killing stationary-phase MRSA for 3 log (**fig. S5J**); ROS-producing agents e.g. hydrogen peroxide (at 220 µM low concentration) synergizes with laser treatment and kills stationary-phase MRSA by 4 log within 2 hours, whereas hydrogen peroxide alone shows minor killing even at 22 mM high concentration (**fig. S5K**). In these cases, besides membrane disruption mechanisms, the depleted antioxidant function of STX contributes to ROS-based killing, consistent with previous findings (*12*).

To determine the clinical relevance of the synergistic therapy between laser treatment and conventional antibiotics, the last-resort antibiotic, daptomycin, was used as the example and further applied on *in vivo* mice skin infection models. To compare the efficacy of different treatment schemes, four groups (control group,10 mg/ml daptomycin-treated group, 10 min laser-treated group, and 10 mg/ml daptomycin plus 10 min laser-treated group) were applied following a 4-day treatment protocol as designed in **fig. S5L**. After the treatment regimen, infected tissue for each mouse was collected with bacterial load quantified via colony-forming unit (CFU) enumeration. The CFU statistical results for each treatment group (**Fig. 5R**) suggest that laser alone-treated group and daptomycin alone-treated group enable 58% and 81% cell killing, respectively; whereas daptomycin plus laser treatment kills around 95% of MRSA in infected skin area. Additionally, the wound areas treated by laser plus daptomycin appear healthier and show the trend of recovery when compared to other groups, as these wound areas show significantly less purulent material, swelling and redness around the edge of the wound. To further evaluate the potential phototoxicity in *in vivo* model, we followed the same treatment protocol as mice skin infection model except removing the MRSA injection step. After the treatment, the skin regions of interest were collected and analyzed via hematoxylin and eosin stained histology slides (representative images shown in **Fig. 5S**). As expected, no phototoxicity induced structure change is observed in the laser-treated group. Additionally, the viability of human epithelial keratinocyte cells is also not affected by laser treatment, even under high laser dosage (**Fig. 5T**). Notably, laser dosage applied in these studies is below the ANSI safety limit for human skin exposure at 460 nm (*23*).

### Inhibition of antibiotic resistance development

To study MRSA response to our phototherapy, we monitored STX expression level during 48-day serial passage study for 10 min laser alone-treated group. Over the course of 48-day passage, steadily decreased STX expression is observed for laser alone-treated group, as resonance Raman peaks for STX drops over serial passage (**Fig. 6A,B**); on 30^th^ and 45^th^ day for two independent replicates, STX abundance drops below the detection limit (**Fig. 6B**); the color of the spun-down cells for both replicates turns to purely white on 48^th^ day whereas the color of the untreated kept golden (**Fig. 6C**). Plate inoculation results further confirm that there is no single colony expressing STX pigment for both replicates after 48-day passage. These results suggest that STX virulence can be eliminated by serial laser treatment without any resistance development. When compared with the original MRSA, the susceptibility of this new phenotype to different antibiotics shows no change or only minor change after serial treatment (**Fig. 6D**).

**Fig. 6.**
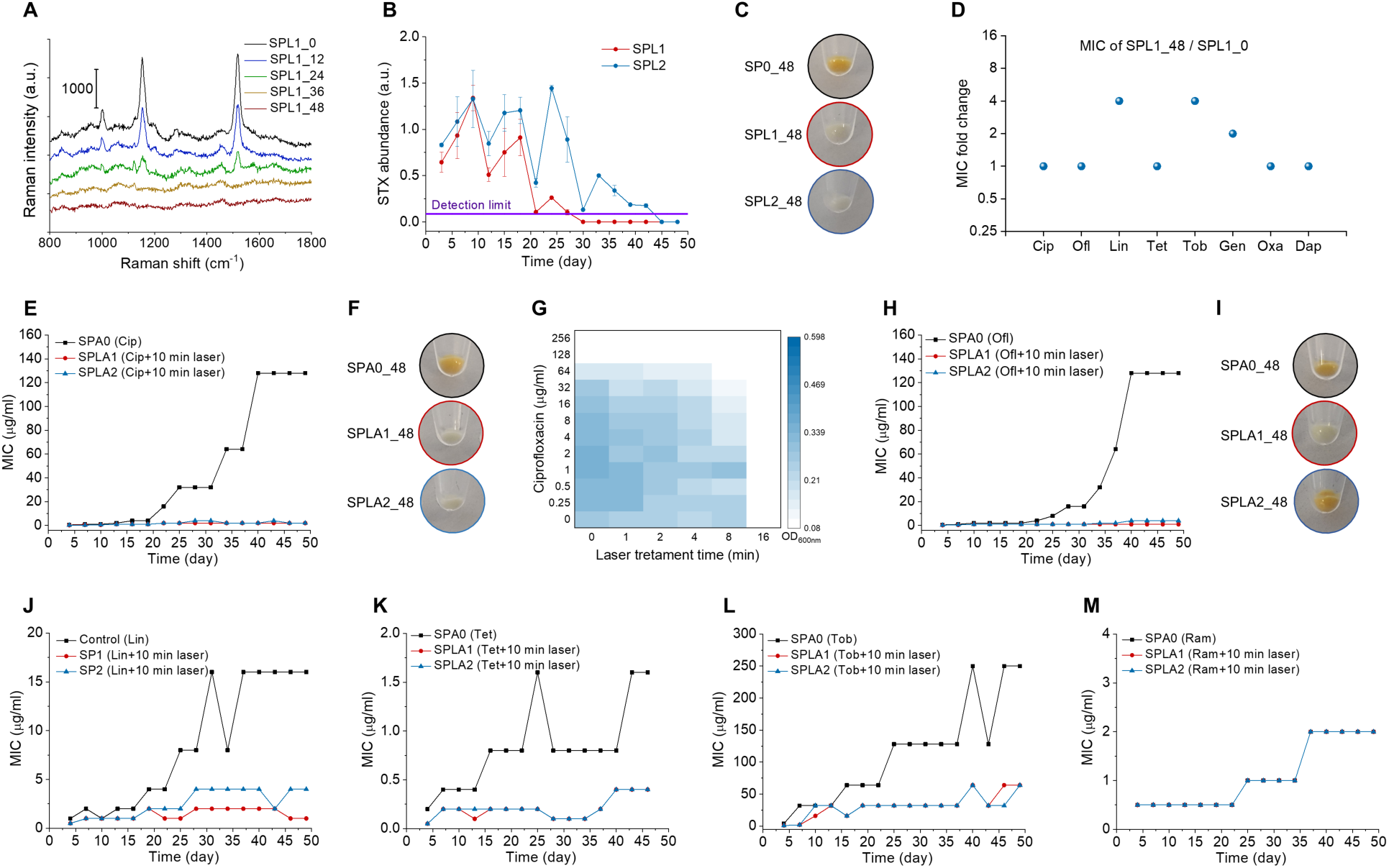
Photo-disassembly of membrane microdomains inhibits resistance development to conventional antibiotics. (A) Representative resonance Raman spectroscopy of STX in stationary-phase MRSA cells at different time checkpoints for the group treated with 10 min laser alone over 48-day serial passage. (B) STX abundance in stationary-phase MRSA cells over 48-day serial passage for groups with or without 10 min laser alone quantified via Raman peak amplitude at 1161 cm^-1^. (C) Images of spun-down cells in (B) after 48-day serially passage showing STX pigmentation. (D) MIC fold change of SPL1 for different classes of antibiotics after 48-day serial passage. (E, H, J, K, L, M) Resistance acquisition over 48-day serial passage in the presence of sub-MIC levels of antibiotics with or without 10 min laser treatment: (E) ciprofloxacin, (H) ofloxacin, (J) linezolid, (K) tetracycline (L) tobramycin, (M) ramoplanin. (F, I) Images of spun-down cells from (E) and (H), respectively, after 48-day serially passage showing STX pigmentation. (G) Checkerboard assay of SPA0_48 showing that 16 min laser treatment completely eliminated cell growth. N=3 for checkerboard assay study, Raman spectra, and STX quantification. SPO, serial passage without any treatment; SPL1 and SPL2, serial passage in independent duplicate with laser treatment alone; SPA0, serial passage with sub-MIC antibiotic treatment alone; SPLA1 and SPLA2, serial passage in independent duplicate with 10 min laser plus sub-MIC antibiotic treatment. The numbers after these abbreviations denote serially passage days.

The development of resistance for different antibiotics with or without 10 min laser treatment was studied in parallel by monitoring MICs for each group in the presence of corresponding antibiotic at sub-MIC level over the course of 48-day passage. Strikingly, with the presence of laser treatment, ciprofloxacin-treated group shows no resistance over the entire passage study, as its MIC is kept ≤ 2 µg/ml, whereas the MIC of ciprofloxacin alone-treated group has reached 128 µg/ml, 256-fold increase relative to its starting MIC (**Fig. 6E**). Its spun-down cells turn purely white for both replicates (**Fig. 6F**); plate inoculation results show that one replicate has no STX expression and the other with mixture of golden and white colonies, consistent with STX expression level monitored through resonance Raman spectroscopy (**fig. S6A**). These results suggest that STX virulence is closely related to ciprofloxacin resistance development via overexpression of efflux pumps (*30*); depletion of STX completely inhibits ciprofloxacin resistance. Therefore, it is highly possible that efflux pump proteins are also co-localized within STX-enriched FMM; STX photolysis malfunctions these efflux pumps while allowing passive diffusion of ciprofloxacin into the cells. Interestingly, using checkerboard assay on the ciprofloxacin-resistant MRSA (MIC: 128 µg/ml), we found that 16 min laser treatment alone completely inhibits the growth of these cells (**Fig. 6G**). This phenomenon suggests that the survival of ciprofloxacin-resistant MRSA relies heavily on STX expression to promote efflux pumps. To further explore this class of antibiotics, ofloxacin was investigated in the serial passage study. Similar results are achieved as shown in **Fig. 6H,I**. After a 48-day serial passage, MIC of ofloxacin alone-treated group reaches 128 µg/ml, whereas ofloxacin plus laser-treated replicates have MICs of 1 and 4 µg/ml, respectively. Based on plate inoculation results, one replicate has pure white colonies and the other had a mixture of white and golden colonies. These results suggest STX photolysis not only increases the susceptibility of MRSA to fluoroquinolones, but also inhibits its resistance development.

Subsequently, laser treatment-mediated resistance inhibition is also found for other antibiotic classes previous found to synergize with STX photolysis, including linezolid, tetracycline, and tobramycin (**Fig. 6J-L**). Delayed resistance development is shown for oxacillin and gentamicin during early serial passages (**fig. S6B,C**). In contrast, decreased resistance development is not shown for ramoplanin, a drug that targets cell wall biosynthesis (**Fig. 6M**), as it is not closely related to the membrane disruption mechanisms. Collectively these results further unveil the causality between STX virulence and antibiotic resistance, as well as demonstrating a way to inhibit resistance development to several major classes of antibiotics via photo-disassembly of FMM.

## Discussion

Current antimicrobial development pipeline has failed to meet the growing needs of new and effective antibiotics to fight bacterial infections (*7*). Here, we demonstrate an unconventional phototherapy approach to combat MRSA antibiotic resistance by targeting its STX virulence factor. This approach fundamentally relies on the interaction between photons and its endogenous chromophores. Although antimicrobial effect of blue light has been documented for decades (*31-33*), the underlying mechanism is still a mystery and its treatment efficacy is limited, hampering its clinical applications. Here, we show that STX is the molecular target of photons and subject to photolysis in the entire blue range. This finding directly challenges the traditionally well accepted hypothesis of blue light-sensitive endogenous porphyrins, meanwhile, profoundly opens new opportunities in this field. The detailed study of STX photochemistry and its photolysis kinetics further suggest a short-pulsed laser to nonlinearly accelerate STX photolysis efficiency, speed, and depth, that are beyond the reach of low-level light sources.

We further show that STX photolysis disorganizes and malfunctions membrane for antibiotic defense in three distinct aspects. First, the disruption renders membrane permeable to antibiotic that target intracellular activities e.g. fluoroquinolones and aminoglycosides. Second, membrane becomes more fluid that facilitates the membrane insertion of membrane targeting antibiotic, e.g. daptomycin. Third, proteins, e.g. PBP2a, that anchors within in the FMM is detached and malfunctioned to defense penicillins. These membrane damage mechanisms demonstrate a novel approach to revive a broad spectrum of conventional antibiotic to combat MRSA. Noteworthy, this approach is fundamentally different from photodynamic therapy (*34*), as it relies on endogenous STX to disrupt cell membrane, thus specifically targeting *S. aureus*, instead of using externally administrated photosensitizer-induced ROS for unselective bacterial eradication.

STX-targeted phototherapy has shown promising potential as a novel treatment platform. Future studies can examine synergies with other classes of antibiotics, as well as the host innate immune system, and/or other reactive oxygen species. For example, disassembly of FMM could be further extended to revive chloramphenicol, as its resistance primarily due to the overexpression of *norA*-encoded multidrug-resistance efflux pumps within the microdomains (*35*). As STX has the antioxidant function to shield MRSA from attacks by ROS, effective STX photolysis could further render MRSA susceptible to oxidative host killing including macrophage cells and neutrophils (*17*). Similar to daptomycin, the modulation on cell membrane fluidity via laser treatment can facilitate non-oxidative host defense of cationic antimicrobial peptides (*25*). Moreover, this platform can be further exploited to screen lead compounds, particularly for those with intracellular targets.

Targeting MRSA STX virulence by photons exemplifies the approach that utilizes the photochemistry between photons and endogenous chromophores to develop a phototherapy platform for bacterial infections. Carotenoids that has structural and functional similarity broadly present in many other bacterial and fungal species (*27*), thus can be photochemically decomposed or modulated in a similar manner. Notably, pigmentation is a hallmark for many pathogenic microbes; these pigments similarly promote microbial virulence and exhibits pro-inflammatory or cytotoxic properties (*36*). Therefore, these pigments could be the targets of photons via either photochemistry or photothermal approach. Several bacterial enzymes that regulate their virulence are also found sensitive to photons (*37*). Therefore, phototherapy approaches based on these specific photon-chromophore interactions could be further explored along this direction.

## Funding

This work was supported by a Keck Foundation Science & Engineering Grant and National Institute of Health Grant R01GM118471 to J.X.C., SPIE-Franz Hillenkamp Postdoctoral Fellowship to J.H. We thank Harvard Center for Biological Imaging for their support for the use of structured illumination microscopy, Boston University Biomedical Engineering Core Facilities for their support for the use of laser scanning confocal microscopies, Boston University School of Medicine Experimental Pathology Laboratory Service Core for analysis and interpretation of mice skin histology.

## Author contributions

J.H., P.T.D. and J.X.C. conceived the project. J.H., P.T.D., L.J.L. designed, organized, performed the studies and analyzed the data. J.H. proposed and demonstrated the short-pulsed laser. J.H. designed, performed and analyzed the Raman measurements. J.H. and P.T.D. designed, performed, and analyzed the membrane disruption fluorescence assays and imaging experiments. J.H., P.T.D. and L.J.L. designed, performed, analyzed the *in vitro* and *in vivo* experiments. J.H. and P.T.D. designed, performed, and analyzed the selection of resistant mutants. C.Q. and T.M. designed, performed and analyzed the molecular dynamics simulations. J.J.L. and P.T.D. performed the western blotting assay. G.Y.L. and E.R.U. provided the bacterial mutants and constructive discussions. M.N.S. provided the clinical bacterial isolates and constructive discussions. Y.W.Z. and S.J. helped in the *in vitro* studies. C.Z. helped in the Raman measurements. J.X.C. supervised the overall project. J.H., P.T.D. and J.X.C. co-wrote the manuscript. All author contributed to discussing and editing the manuscript.

## Competing interests

Authors declare no competing interests.

## Data and materials availability

All data needed to evaluate the conclusions in this study are present in the main manuscript and/or the Supplementary Information. Additional data related to this study may be requested from the authors.

## Supplementary Materials

Materials and Methods

Figures S1-S6

Tables S1

