## Supplementary Materials for "Photo-Disassembly of Membrane Microdomains Revives Conventional Antibiotics against MRSA"

**This PDF file includes:**

Materials and Methods
Figs. S1 to S6
Tables S1

### Materials and Methods

Nanosecond pulsed laser and LED systems

The nanosecond pulsed laser system was composed of a nanosecond pulsed laser source (Opolette HE355 LD, OPOTEK Inc.), a 1 mm-core multimode fiber for light delivery (NA=0.22, OPOTEK Inc.), and a custom-built handheld device. Key specifications of the laser source: tunable wavelength range, 410-2400 nm; pulse repetition rate, 20 Hz; maximum pulse energy at 460 nm, 8 mJ; pulse duration, 5 nanoseconds (ns); spectral linewidth, 4-6 $cm^{-1}$; pulse-pulse stability, <5%. Within the handheld device, a collimation lens (LB1471-A, Thorlabs) was applied to expand the output beam with a diameter of 1 cm. This device was mounted on a stable optical table for experiments shown in Fig. 1a. After collimation by all these optical components, this system provides a final maximum output of 120 mW (6 mJ in pulse energy). Within the illumination area, photon density follows a near-Gaussian distribution. With the diameter of sample droplet at around 5 mm, the photon density over the sample droplet in this study was assumed uniform.

The continuous-wave LED system applied in this study was composed of a blue light LED (M470L3, Thorlabs), an adjustable collimation adapter (SM2F32-A, Thorlabs), and a power controller (LEDD1B, Thorlabs). The output of the blue light LED is centered at 465 nm with bandwidth of 25 nm and maximum power of 650 mW. The output power of the LED system was adjustable, and its beam size was controlled through the collimator and an iris. In order to compare with nanosecond pulsed laser, the output power of the LED was set to 120 mW and used to illuminate an area of 1 cm in diameter.

Bacterial strains and growth conditions

Methicillin-resistant *S. aureus* (MRSA USA 300, NRS 384), *S. aureus* ΔCrtM mutant, vancomycin-resistant *S. aureus* (VRSA 9, NR-46419), Methicillin-resistant *S. aureus* (MRSA USA 500, NRS 385).

Log-phase and stationary-phase bacterial inoculum preparation: colonies from streaked plate of frozen bacterial stock were inoculated in sterile tryptic soy broth (TSB, 22092, Sigma Aldrich) medium and grown in an orbital incubator (12960-946, VWR) with a shaking speed of 200 rpm for 2-3 hours at 37 ℃for log-phase bacteria (~$10^7$ cells/ml). Before each experiment, bacterial cells were spun down and then the harvested bacteria pellets were washed with 1×phosphate-buffered saline (PBS) twice and then resuspended in 1×PBS at its original concentration. Stationary-phases bacterial solution were prepared following the same procedure except that bacteria inoculum was cultured to three days.

Antibiotics and chemicals

Antibiotics used in this study: daptomycin (103060-53-3, Acros Organics), oxacillin (28221, Sigma Aldrich), gentamicin (G1914, Sigma Aldrich), tobramycin (T4014, Sigma Aldrich), ciprofloxacin (17850, Sigma Aldrich), ofloxacin (O8757, Sigma Aldrich), linezolid (PZ0014, Sigma Aldrich), tetracycline (87128, Sigma Aldrich), ramoplanin (R1781, Sigma Aldrich), vancomycin (V2002, Sigma Aldrich). 10 mg/ml stocks of all compounds were made in 1×PBS or DMSO (W387520, Sigma Aldrich) or sterile water. For treatments with daptomycin, sterile medium or buffer was supplemented with $CaCl_2$ (C79-500, Fisher Scientific) with final working concentration of 50 μg/ml.

Fluorescent dyes used in this study: SYTOX green (S7020, Thermo Fisher Scientific), Texas red-X, succinimidyl ester, single isomer (T20175, Thermo Fisher Scientific), FITC-dextran (FD4, FD70, FD500, Sigma Aldrich). $DiIC_{18}$ (1,1'-Dioctadecyl-3,3,3',3'-Tetramethylindocarbocyanine Perchlorate, D282, Thermo Fisher Scientific). BODIPY FL, STP ester, sodium salt (B10006, Thermo Fisher Scientific).

Resonance Raman spectroscopy

STX was quantified by its Raman peak amplitude at 1161 $cm^{-1}$ measured by resonance Raman spectroscopy (1221, LABRAM HR EVO, Horiba) with a 40×objective (Olympus) and an excitation wavelength of 532 nm. Samples (either from bacterial colony or STX extract solution) were sandwiched between two glass cover slides (48393-230, VWR international) with a spatial distance of ~80 µm. To study staphyloxanthin photolysis kinetics, the same samples were measured after each laser treatment.

Staphyloxanthin extraction protocol

The STX extraction protocol was adapted from a previous report (*1*). Briefly, 2 ml of stationary-phase MRSA were spun down, washed with 1×PBS. Then the MRSA pellets were harvested through centrifuge and crude STX pigment was extracted by 200 µl warm methanol in dark at 55 °C for 20 min.

Absorption spectroscopy

Absorption spectroscopy of MRSA solution was performed after different laser treatment time. Briefly, stationary-phase MRSA stationary-phase MRSA (~$10^8$ cells/ml) was washed and suspended into 1×PBS at its original concentration. Aliquots of 100 µl was transferred into a 96 well plate. The absorption spectrum of the lidded wells after each laser treatment (1.5 min laser treatment interval) were monitored by a plate reader (SpectraMax i3x, Molecular Devices) with a spectral window of 300-800 nm and a step size of 2 nm. For the treatment, each well was directly illuminated by laser beam from the well top (1 cm diameter illumination area, 120 mW). Three independent replicates were applied in the study.

Fluorescence microscopic imaging techniques

For super-resolution imaging, we used a structured illumination microscope (ELYRA super-resolution microscope, Zeiss) with a 100×oil objective. There are several diode lasers used as the excitation sources in the system (405 nm, 488 nm, 561 nm, 638 nm). In the case of FITC-dextran and PBP2a immunofluorescence imaging, we used excitation wavelengths of 488 nm and 561 nm, respectively. Image processing and analysis were directly performed with the provided software for the system.

For the confocal laser scanning microscope, we used a laser scanning confocal microscope (FV3000, Olympus) with two high-sensitivity GaAsP/GaAs photomultiplier tubes (PMTs). The images demonstrated in this study were acquired in a high-speed resonant Galvo-Galvo scanning mode and via an UPLSAPO 100×oil objective (NA=1.35, Si oil immersion, 0.2 mm working distance). Inside this confocal microscope, there are six solid state diode lasers (405 nm, 445 nm, 488 nm, 514 nm, 561 nm, 640 nm). In the case of SYTOX green, FITC-dextran dyes, and daptomycin-BODIPY, we used an excitation wavelength of 488 nm. For the $DiIC_{18}$, we used 561 nm as the excitation wavelength. Gentamicin-Texas red was excited by a 514-nm laser.

SYTOX green membrane permeability assay

Briefly, 1 ml of stationary-phase MRSA (~$10^8$ cells/ml) was spun down, got rid of the supernatant, and resuspended with 100 µl of sterile 1×PBS. 5 µl of the above solution was then exposed into laser beam with different treatment time (laser power, 120 mW; illumination area, 1 cm in diameter). After treatment, MRSA solution was collected into 985 µl of sterile water, as SYTOX green shows best performance in buffers without phosphate. Subsequently, 10 µl of stock SYTOX green solution (5 mM in DMSO) was supplemented before aliquoting into a 96-well plate. The fluorescence emission intensity at 525 nm (excitation at 488 nm) was monitored by a plate reader (SpectraMax i3x, Molecular Devices) for more than 2 hours with a 5-min interval at room temperature. To further visualize the uptake of SYTOX green under a laser scanning confocal fluorescence microscopy, MRSA cells were further prepared following these steps: spin down MRSA pellets, get rid of the supernatant, wash the pellets with sterile water twice, and fix them with 10% formalin (HT501128-4L, Sigma Aldrich). All experiments were conducted in duplicate or triplicate.

FITC-dextran membrane permeabilization assay

To estimate how large a molecule can diffuse into the damaged membrane, we applied dextran conjugated with fluorescein isothiocyanate (FITC-dextran) with variable molecular weight/Stokes radius (FD4-FD500, Sigma Aldrich) and monitored their insertion before and after laser treatment. Briefly, 1 ml of stationary-phase MRSA (~$10^8$ cells/ml) was spun down, got rid of the supernatant, and resuspended with 100 µl of sterile 1×PBS. 5 µl of the above solution was exposed to pulsed laser with different treatment time. After laser treatment, bacterial solution was collected into 985 µl of sterile pre-warmed TSB, supplemented with 10 µl of FITC-dextran (1 mg/ml), and incubated for 30 min at 37°C. The integrated fluorescence signal from an aliquot of the bacterial solution with or without laser treatment was measured through a plate reader with excitation of 488 nm and emission of 520 nm, respectively. Meanwhile, after incubation, the bacterial solution was spun down, got rid of the supernatant, washed with pre-warmed TSB twice, and fixed with 10% formalin. Structured illumination microscopy was conducted to quantify FITC-dextran uptake and its distribution on cell membrane with an excitation wavelength of 488 nm. Quantitative analysis of fluorescence emission intensity from individual MRSA cells was performed among groups with different laser treatment time.

Gentamicin-Texas red intracellular uptake assay

To study laser-mediated intracellular uptake of gentamicin (a representative of aminoglycoside), gentamicin was conjugated with a fluorescent dye, Texas-red, to form gentamicin-Texas red. Briefly, 10 mg of gentamicin was dissolved into 1 ml of 0.1 M sodium bicarbonate buffer (S8761-500ML, Sigma Aldrich). 10 mg/ml of Texas red-X succinimidyl ester (T6134, Thermo Fisher Scientific) was added to the gentamicin solution slowly drop by drop. Then the mixed solution was stirred at room temperature for 1 hour. Gentamicin-Texas red was purified through sufficient dialysis against 0.1 M sodium bicarbonate buffer in a dialysis sack (Slide-A-Lyzer G2 Dialysis Cassettes, 2K MWCo, 15 mL, 87719, Thermao Fisher Scientific), and harvested through lyophilization (Labconco). Next, 1 ml of stationary-phase MRSA (~$10^8$ cells/ml) was spun down, got rid of the supernatant, and suspended with 100 µl of sterile 1×PBS. 5 µl of the above solution was exposed to pulsed laser for different treatment time (1 cm diameter illumination area, 120 mW). After treatment, bacterial droplet was collected into 985 µl of sterile 1×PBS, and then add 10 µl of 1 mg/ml Gentamicin-Texas red. Mixed solution was incubated at 37°C for 30 min with a shaking speed of 200 rpm. After incubation, MRSA pellets were harvested through washing with sterile 1×PBS twice and then fixed with 10% formalin. Visualization of gentamicin-Texas red on bacterial cells was achieved through a confocal laser scanning microscope (FV 3000, Olympus) with the excitation wavelength of 514 nm. Quantitative analysis of fluorescence emission intensity from individual MRSA cells was conducted and allocated among groups with different laser treatment time.

Ciprofloxacin intracellular uptake assay

To understand how laser treatment affects the uptake of ciprofloxacin (a representative of fluoroquinolone), we adopted a protocol published elsewhere (*2*). Briefly, 1 ml of stationary-phase MRSA (~$10^8$ cells/ml) was spun down, got rid of the supernatant, and suspended with 100 µl of sterile 1×PBS. 5 µl of the above solution was exposed to pulsed laser for different treatment time (1 cm diameter illumination area, 120 mW). After treatment, bacterial droplet was collected into 994 µl of sterile 1×PBS, and then added 1 µl of 10 mg/ml of ciprofloxacin (17850-5G-F, Sigma Aldrich), then incubated for 30 min at 37°C with a shaking speed of 200 rpm. After incubation, MRSA pellets were washed twice by 2 ml of ice-cold PBS. Then ciprofloxacin was extracted using 1 ml of glycine (G8898, Sigma Aldrich)-HCl buffer at pH=3 for 2 hours. The amount of ciprofloxacin was estimated and quantified by measuring the fluorescence intensity via a plate reader with an excited wavelength of 275 nm and emission wavelength of 410 nm.

$DiIC_{18}$ membrane fluidity assay

$DiIC_{18}$ is a fluorescent dye that displays affinity for membrane areas with increased fluidity due to its short hydrocarbon tail (*3*). In our protocol, briefly, 1 ml of stationary-phase MRSA (~$10^8$ cells/ml) were spun down, got rid of the supernatant, and suspended with 100 µl of pre-warmed TSB supplemented with 1% DMSO. 5 µl of the above solution was exposed to pulsed laser for different treatment time. After treatment, bacterial droplets (with 2.5, 5, 10 min treatment time) were collected into 985 µl of pre-warmed TSB supplemented with 1% DMSO. 10 µl of $DiIC_{18}$ (stock:10 mg/ml in DMSO) were added to the above solution, and incubated for 30 min at 37°C. After incubation, harvested MRSA pellets were washed with pre-warmed TSB supplemented with 1% DMSO for four times, then sandwiched the concentrated bacterial samples between a poly-prep cover slides (P0425, Sigma Aldrich) and a thin cover glass (48404-457, VWR international). A confocal laser scanning microscope (FV3000, Olympus) was applied to visualize and quantify $DiIC_{18}$ uptake at an excitation wavelength of 561nm and via a 100×oil immersion objective (NA = 1.35, Olympus).

Daptomycin-BODIPY membrane insertion assay

To study how the membrane fluidity change affects the insertion of membrane-targeting antibiotics, we applied daptomycin-BODIPY membrane insertion assay detailed as below. Firstly, we conjugated daptomycin with a fluorescent dye, BODIPY STP ester (B10006, Thermo Fisher Scientific). Briefly, 10 mg of daptomycin (103060-53-3, Acros Organics) was dissolved into 1 ml of 0.1 M sodium bicarbonate solution. Then 100 µl of BODIPY STP ester (B10006, Thermo Fisher Scientific, stock: 1 mg/ml in DMSO) was added to the daptomycin solution drop by drop. Then the mixed solution reacted under stirring at room temperature for 1 hour. Afterwards, the solution was under overnight dialysis against extensive 0.1 M sodium bicarbonate solution. After dialysis, the mixed solution was lyophilized. To further label MRSA cell membrane with daptomycin-BODIPY, 1 ml of stationary-phase MRSA (~$10^8$ cells/ml) was spun down, got rid of the supernatant, and suspended with 100 µl of sterile 1×PBS. 5 µl of the above solution was exposed to pulsed laser with different treatment time. After treatment, bacterial droplets were collected into 985 µl of sterile pre-warned TSB medium containing 150 µg/ml of $CaCl_2$. 10 µl of daptomycin-BODIPY (stock: 3 mg/ml in 1×PBS) was added to the above solution, and incubated for 30 min at 37°C. After incubation, harvested MRSA pellets were washed with 1×PBS twice, and fixed with 10% formalin. Confocal laser scanning microscope (FV3000, Olympus) was conducted to quantify daptomycin-BODIPY distribution and its signal intensity at an excitation wavelength of 488 nm. Quantitative analysis of the signal from individual MRSA cells was performed among groups with different laser treatment time.

PBP2a immunofluorescence assay

Basically, 1 ml of stationary-phase MRSA (~$10^8$ cells/ml) was spun down, got rid of the supernatant, and suspended with 100 µl of sterile 1×PBS. 5 µl of the above solution was exposed to pulsed laser for different treatment time. After treatment, bacterial droplets were collected into 980 µl of sterile 1×PBS, and 20 µl of a primary antibody (Rabbit Anti-PBP2a, RayBiotech, 130-10073-20, 10 µg/ml) targeting PBP2a was added to the above solution. Then the mixed solution was incubated for 30 min at 37°C with a shaking speed of 200 rpm. After incubation, MRSA pellets were washed twice with sterile 1×PBS. As the last wash, MRSA pellets were suspended with 990 µl of 1×PBS. Then 10 µl of secondary antibody (Goat anti-Rabbit Cy5, Abcam, ab97077, 0.5 mg/ml) was added to the above solution, incubated for another 30 min at 37°C with a shaking speed of 200 rpm. After incubation, MRSA pellets were washed with sterile 1×PBS twice and fixed with 10% formalin. Immunofluorescence experiment was conducted by a confocal laser scanning microscope at an excitation wavelength of 650 nm. Quantitative analysis of signal intensity and its distribution from individual MRSA cells was performed among groups with different laser treatment time.

PBP2a western blotting assay

Briefly, 3 ml of stationary-phase MRSA (~$10^8$ cells/ml) was spun down and suspended with 100 µl of 1×PBS. 20 µl of the mixed solution was aliquoted to a centrifuge tube (89166-280, VWR international), and then exposed to pulsed laser with different treatment time (control, 5 min, 10 min, 20 min). After exposure, the four tubes containing MRSA solution were spun down at a speed of 13,000×g for 10 min at 4°C. Then the supernatants were collected into four new sterile tubes. To extract proteins from MRSA pellets, after removing the supernatant, MRSA pellets were suspended with 100 µl of lysis buffer (96.8 µl of RIPA, 1 µl 500 mM DTT, 1 µl of 10% Triton-X, 1 µl of protease inhibitor, and 1 µl of phosphorylase inhibitor). Then the mixed solutions were sonicated by a sonication probe (Cole-Parmer) at 4°C. Released proteins were harvested from the supernatants by centrifuging at 13,000×g for 10 min at 4°C. Electrophoresis separation of the proteins from both MRSA pellets and supernatants was conducted on a 12% SDS-PAGE gel (stacking gel: 4%) at a voltage of 50 V for 30 min followed by 100 V for 1 hour in 1×running buffer (1610772, Bio-Rad). After separation of the proteins, gels were transferred to a PVDF membrane (1620184, Bio-Rad) at a current of 150 mA overnight at 4°C in 1×transfer buffer (1610771, Bio-Rad). After transferring, PVDF membrane was harvested and put into a clean plastic reservoir containing 5% milk solution (1706404, Bio-Rad). Then the plastic reservoir was placed on a rocking shaker for 30 min. After blocking, the PVDF membrane was further labelled with primary antibody (Rabbit anti-PBP2a, 1:500 dilution in 5% milk solution) for 2 hours in a rotary shaker. Then the PVDF membrane was washed with 1×washing buffer three times with each time for 5 min on the rotary shaker. Afterwards, the PVDF membrane was conjugated with a fluorescent secondary antibody (*Eu*-anti-Rabbit, Molecular Devices, 1:1000 dilution in 5% milk solution) for 1 hour on the rotary shaker and then washed with 1×washing buffer three times with each time for 5 min on the rotary shaker. Lastly, the protein-antibody-antibody fluorophore complex was detected through a plate reader at an excitation wavelength of 340 nm.

Membrane computational method

The Coarse-Grained (CG) simulations were performed using the MARTINI forcefield. The parameters for the cardiolipin were taken from the MARTINI database (*4*). For PBP2a, only the transmembrane helix was included in this simulation as we focus on the membrane properties in the current work. The saturated and unsaturated tail of the STX lipid were modeled by "C1" and "C4" bead type, respectively following other lipid parameters within the MARTINI model. The head group of the STX lipid is a glucose for which the MARTINI parameters were taken from the database. The bond and angle parameters for the CG beads of the STX tails were determined using structural information obtained from atomistic simulations. A single STX lipid in solution was simulated using the all atom CHARMM27 (*5*) forcefield and the TIP3P (*6*) water model. The equilibrium bond length and angle for the STX tail CG beads were obtained from the positions of the mass centers of the corresponding groups in atomistic simulations. The bond force constants for both the saturated and unsaturated tails and the angle force constants for the saturated tail were taken as same as for the other lipids in the MARTINI model. However, since every other bond in the unsaturated tail is a C=C bond, the tail is expected to be very rigid. So, the angle force constants for the unsaturated tail were taken to be higher ($200\ kJ/mol - rad^2$) than the angle force constants for the saturated tail ($25\ kJ/mol - rad^2$). To model STX following its photolysis, the long unsaturated tail was truncated, as suggested by the complete loss of C=C vibrational peak in the Raman spectra after STX photolysis. The transmembrane helix of the PBP2a protein was generated using the Chimera software (*7*). The CG parameters for the peptide were generated using a script provided in the MARTINI database. We built a bilayer (~17×17 $nm^2$) of randomly mixed STX, cardiolipin and peptides (400:200:36). The built system was then solvated using the MARTINI water model; 10% anti-freezing beads were also added to avoid any artificial water freezing. Sodium and chlorine ions were then added to maintain 150 mM salt concentration. Each system (with full and truncated STX lipids, respectively) was equilibrated and simulated under the constant pressure and constant temperature ensemble for 10 μs. All simulations were conducted using the GROMACS program (*8*).

The RDF or the pair correlation function, $g(r)$, between molecule type $A$ and molecule type $B$ is calculated using the following equation

$$g(r) = \frac{1}{<\rho_B>} \frac{1}{N_A} \sum_{i}^{N_A} \sum_{j}^{N_B} \frac{\delta(r_{ij} - r)}{4\pi r^2}$$

Here, $N_A$ and $N_B$ are the number of molecules of type $A$ and type $B$, respectively. $\rho_B$ denotes the density of molecule type $B$ in a sphere of radius $r_m$ around the molecule type $A$ and $< \rho_B >$ is the average of $\rho_B$ calculated over all type $A$ molecules. The $r_m$ was taken to be ~6 nm which is half of the shortest box dimension.

The area expansion modulus $K_A$ of the membrane was calculated using the following equation:

$$K_A = \frac{k_B T < A >}{< \delta A^2 >}$$

Here $k_B$, $T$, and $A$ are the Boltzmann constant, absolute temperature and the membrane surface area, respectively; $< \delta A^2 >$ represents the fluctuation in the surface area, which was calculated as $< \delta A^2 > = < (A - < A >)^2 >$, where $< A >$ is the mean value of the surface area averaged over ~5 µs simulation. The thermal fluctuations in the membrane surface area is less in a tightly packed membrane. Thus, a higher value of $K_A$ represents a more tightly packed membrane.

Bacterial growth kinetics

To monitor the response of bacteria to laser treatment alone, antibiotic treatment alone, or their combinations, bacterial growth was continuously monitored overnight (18 hours with an interval of 30 min at 37°C) by measuring optical density at 600 nm ($OD_{600}$). Depending on the specific assay applied, the bacterial cells were suspended in 100 or 200 µl TSB medium under different treatment schemes (antibiotic alone, laser treatment alone, antibiotic plus laser treatment) with a final concentration of ~$10^5$ CFU/ml. Bacterial growth was defined as $OD_{600} \geq 0.1$.

Minimal inhibitory concentration measurement

The MICs of antibiotics were determined by the standard broth-dilution method recommended by the Clinical and Laboratory Standards Institute (*9*). Briefly, bacterial strains were grown aerobically overnight on tryptic soy agar (TSA, 22091, Sigma Aldrich) plates at 37°C. Bacterial colonies were then suspended into TSB medium with a concentration of ~$10^5$ CFU/ml and then transferred into 96-well plates (71000-078, VWR international). Antibiotics were added in the first row of the 96-well plates and then two-fold serially diluted. Plates were then incubated aerobically at 37°C for ~18 hours. MICs reported were the minimum concentration of antibiotics that completely inhibited the visual growth of the bacteria or with $OD_{600}$ less than 0.1 monitored by a plate reader (SpectraMax i3x, Molecular Devices). For each measurement, three independent replicates were applied. Supplementary Table 1 shows the MICs of selected antibiotics against the tested bacterial strains.

Colony-forming-unit enumeration assay

To quantify viable bacterial cells, CFU experiments were performed. 100 µl of sample analyte was transferred into a 96-well plate and then three or four ten-fold serial dilution achieved by transferring 20 µl bacterial culture into 180 µl 1×PBS in the next dilution row. After serial dilution, an aliquot (4 µl) from each well was spotted onto sterile TSA plates. After incubating the plates overnight (~18 hours) at 37 °C, the colonies were enumerated, and cell number was calculated in CFU/ml. For each CFU enumeration experiment, three independent replicates were applied.

Post-exposure and post-antibiotic assays

To study the post-exposure effect for laser treatment, stationary-phase MRSA was prepared, washed and resuspended in 1×PBS at its original concentration. An aliquot (5 µl) of the bacterial suspension was transferred onto a glass cover slide (48393-230, VWR international) and treated by pulsed laser for different treatment time (1 cm-diameter illumination area, 100 mW). After treatment, the droplets were collected and resuspended 1:1000 into 5 ml of TSB medium for each group. An aliquot of 100 µl was then transferred to a 96-well plate for growth monitoring.

To study the post-antibiotic effect of antibiotics, we adopted a protocol published elsewhere (*10*). Briefly, stationary-phase MRSA (~$10^8$ cells/ml) were prepared, washed and cultured in fresh TSB at its original concentration supplemented with 4×MIC of antibiotics including ofloxacin, oxacillin and gentamicin for one hour at 37°C. A tube containing the untreated bacterial cells served as a control. Afterwards, antibiotics were washed out and 1:1000 diluted in TSB. An aliquot of 100 µl was then transferred to a 96-well plate for growth monitoring. Three independent replicates were applied for each antibiotic and/or laser-treated groups. Post-antibiotic effect was estimated by the difference between the times that required for both the control and antibiotic-treated groups to reach $OD_{600} = 0.3$.

Checkerboard broth dilution assay

Stationary-phase bacterial cells was washed and resuspended in 1×PBS at its original concentration. An aliquot (5 µl) of the bacterial solution (used as a control group) was transferred onto a glass cover slide as a droplet of ~5 mm in diameter and exposed to pulsed laser for different treatment time (1-cm diameter illumination area, 100 mW). The treated droplet was collected and resuspended into 5 ml TSB (1:1000 dilution) for each group. Corresponding groups without laser treatment were also conducted for comparison. The bacterial suspensions were transferred to a 96-well plate with antibiotics supplemented into with the first row of the 96-well plate for eight two-fold serial dilution starting at a desired antibiotic concentration (e.g. ofloxacin: 2 µg/ml). After serial dilution, bacterial growth within the same well plate was monitored by a plate reader for 18 hours ($OD_{600}$, 37 °C). The checkerboard assay was used for groups with laser treatment alone or laser plus antibiotic treatment. Two independent experiments of checkerboard assay were performed for each antibiotic with or without laser treatment. Based on the readout of $OD_{600}$, a heat map was created to evaluate the antibiotic potentiation or synergistic effect enabled by STX photolysis.

Synergy evaluation between antibiotic and laser treatment

Based on the checkerboard results, the fractional inhibitory concentration index (FICI), a synergy evaluation method between two antibiotics, was calculated as below: FICI = MIC of antibiotic A in combination / MIC of antibiotic A alone + MIC of antibiotic B in combination / MIC of antibiotic B alone. The interaction of the two antibiotics was defined as below: synergy if FICI ≤ 0.5, no interaction if 0.5 < FICI ≤ 4, antagonism if FICI > 4 (*11*). As this demonstrated phototherapy approach depletes STX virulence instead of completely inhibiting bacterial growth, thus there is no MIC for laser treatment alone. Considering this reason, the synergy calculation was simplified as below: FICI = MIC of antibiotic A in combination with laser treatment / MIC of antibiotic A alone with synergy defined by FICI ≤ 0.5.

Time-killing assay

Stationary-phase MRSA was prepared, washed and resuspended in 1×PBS at two-times of its original concentration. An aliquot (5 µl) of the MRSA suspension was transferred onto a glass cover slide as a droplet of ~5 mm in diameter and exposed to pulsed laser for different treatment time (1 cm diameter illumination area, 100 mW). After laser treatment, the droplets were resuspended into 200 µl of 1×PBS (1:40 dilution) supplemented with antibiotics at different concentrations in a mini centrifuge tube (89166-278, VWR international). For example, daptomycin was added into MRSA solution after laser treatment at desired concentration of 0×MIC, 5×MIC, 10×MIC, 30×MIC, or 100×MIC supplemented with 50 µg/ml $CaCl_2$). Corresponding groups without laser treatment were also conducted for comparison. These tubes were incubated within an orbital incubator (37 °C, 200 rpm) for different incubation time. At each specific time point, 40 µl of aliquot from each group was transferred to a 96-well plate for follow-up CFU enumerating assay. In the case of tobramycin, additional antibiotic washing by 1×PBS was performed before the CFU experiment to avoid antibiotic interference. For time-killing assay in fresh human whole blood, similar protocol was followed as above, except that 1×PBS was replaced by fresh human whole blood and the initial stationary-phase MRSA solution was diluted by ten times with a concentration of ~$10^7$ CFU/ml. The time-killing assay for hydrogen peroxide also followed the same protocol except replacing supplemented antibiotic by low-concentration hydrogen peroxide.

Serial passage assay for resistance development

To understand whether laser treatment could cause genotypic or phenotypic change in MRSA, and whether STX photolysis could reduce the resistance development for conventional antibiotics, a serial passage study for each treatment scenario was conducted. The initial generation (Day 1) used in this study was stationary-phase MRSA. The sample was prepared, washed and resuspended in 1×PBS at its original concentration. An aliquot (5 µl) of the MRSA suspension was transferred onto a glass cover slide as a droplet of ~5 mm in diameter with or without 10 min laser treatment (1 cm diameter illumination area, 120 mW). The droplets were then collected and resuspended into 5 ml of TSB medium (1:1000 dilution) with an estimated cell concentration of $10^5$ CFU/ml. To study resistance development or selection induced by laser treatment alone, three groups were included: a group without laser treatment (SP0), a group with laser treatment (SPL1), and another independent group with laser treatment as a duplicate (SPL2). To study resistance development induced by antibiotic treatment alone and laser plus antibiotic treatment, three groups were included for each antibiotic: antibiotic alone-treated group (SPA0), laser plus antibiotic-treated group (SPLA1), and another laser plus antibiotic-treated group as another independent serial passage (SPLA2). For SP0, SPL1, and SPL2, 200 µl of bacterial suspension was directly transferred to each well of a 96-well plate, with three replicates conducted for each group. For SPA0, SPLA1, and SPLA2, 200 µl of bacterial suspension was transferred into the first dilution row of a 96-well plate with supplemented antibiotics at a desired starting concentration, whereas 100 µl of bacterial suspension was transferred to the rest dilution rows. After twelve two-fold serial dilution, 100 µl of bacterial suspension was added into each well to make a 200 µl of final volume for each well, thus, as an example, supplementing 5.12 µl of 10 mg/ml ofloxacin solution into 200 µl of bacterial culture in the first dilution row makes a starting concentration of 128 µg/ml. Three replicates were applied for each group. These well plates were incubated in a shaker at 37 °C and 200 rpm for 18 hours followed by $OD_{600}$ measurement by a plate reader. After MICs recording for each group, the well plates were continuously incubated in the shaker for 3 days in total. On Day 4, 200 µl of bacterial sample from each group was collected, washed, and resuspended in 1×PBS at its original concentration used as new inoculum for the next passage following the same protocol as described above. Samples for SPA0, SPLA1, and SPLA2 groups were collected from wells supplemented with sub-MIC antibiotic. Samples for SP0, SPL1, and SPL2 groups were also collected from the well plates. The left bacterial suspension for each group was stored in 25% glycerol at -80 °C for subsequent analysis and experiments. Serial passage for all groups were performed for 50 days with 16 generations in total.

Raman spectroscopy was then applied to monitor STX expression level in groups of interest after the entire serial passage experiment. The protocol is detailed as below: 100 µl of ~400 µl stored bacterial culture was collected, spun down with the supernatant being removed, then resuspended into 5 µl 1×PBS as high-concentration bacterial solution (20 times concentrated). An aliquot (1 µl) was transferred and then sandwiched between two glass cover slides for STX quantification by resonance Raman spectroscopy.

<u>*In vivo* mice infection model</u>

The *in vivo* mice experiment was conducted following protocols approved by Boston University Animal Care and Use Committee (BUACUC). To initiate the formation of a skin wound, five groups (N=5) of eight-week-old female BALB/c mice (obtained from the Jackson Laboratory, ME, USA) were disinfected with ethanol (70%) and shaved on the middle of their back (approximately a one-inch by one-inch square region around the injection site) one day prior to infection as described from a reported procedure (*10*). To prepare the bacterial inoculum, an aliquot of overnight culture of MRSA USA300 was transferred to fresh TSB and shaken at 37 °C until an $OD_{600}$ value of ~1.0 was achieved. The cells were centrifuged, washed once with 1×PBS, re-centrifuged, and then re-suspended in 1×PBS. Mice subsequently received an intradermal injection (40 µl) containing ~$10^9$ CFU/ml MRSA USA300. An open wound formed at the site of injection for each mouse, ~48 hours post-infection. Topical treatment was initiated subsequently with each group of mice receiving the following: daptomycin (1%, using glycerol as the vehicle), pulsed laser (1 cm diameter illumination area, 10 min treatment time, 120 mW), or a combination of pulsed laser and daptomycin. One group of mice was left as the control. Each group of mice receiving a particular treatment regimen was housed separately in a ventilated cage with appropriate bedding, food, and water. Mice were checked twice daily during infection and treatment to ensure no adverse reactions were observed. Mice were treated once daily (once every 24 hours) for three days, before they were humanely euthanized via $CO_2$ asphyxiation 12 hours after the last dose was administered. The region around the skin wound was lightly swabbed with ethanol (70%) and excised. The tissue was subsequently homogenized in 1×PBS. The homogenized tissue was then serially diluted in 1×PBS before plating onto mannitol salt agar plates (*S. aureus* specific). Plates were incubated for at least 19 hours at 37 °C before CFU assay for each group. Outlier was removed based upon the Dixon Q Test. Data were analyzed via an unpaired *t*-test, utilizing Origin 2019b (OriginLab Corporation).

<u>H&E histology analysis of mice skin</u>

To evaluated phototoxicity of laser treatment on healthy mice skin, mice (N=3) were treated with pulsed laser once daily for three days. After treatments, mice were humanely euthanized under $CO_2$ asphyxiation. Treated mice skin were sacrificed and collected into 10% formalin solution. H&E (hematoxylin and eosin) staining were utilized to stain sacrificed mice skin. Skin slices were imaged and analyzed by Boston University Experimental Pathology Service Core.

Phototoxicity on human cell line

To evaluate the toxicity of pulsed laser, we chose a human cell line (human epithelial keratinocyte cells, HEK 293) to evaluate the phototoxicity. HEK cells were cultured at Dulbecco's Modified Eagle Medium (DMEM, Thermo Fisher Scientific) supplemented with 10% fetal bovine serum. A colorimetric MTT assay was used to assess the cell metabolic activity. Briefly, 5 mg of MTT (M6494, Thermo Fisher Scientific) was dissolved in 1 ml 1×PBS. Then MTT solution was diluted with serum-free DMEM medium at ratio of 1:10. The pulsed laser was applied to treat HEK 293 cells in a 96-well plate. After treatment, 100 µl of the diluted MTT solution (pre-warmed) was added to each treated well, and then incubated for four hours in dark at 4°C. After incubation, supernatants were removed, and 200 µl of DMSO was added to the wells. $OD_{540}$ from each treated well was measured by a plate reader.

Statistical analysis

Statistical analysis was conducted through unpaired *t*-test. **** means significantly different with the $p$-value < 0.0001. *** means significantly different with the $p$-value < 0.001. ** means significantly different with the $p$-value < 0.01. * means significantly different with the $p$-value < 0.05. 'ns' means no significant difference.

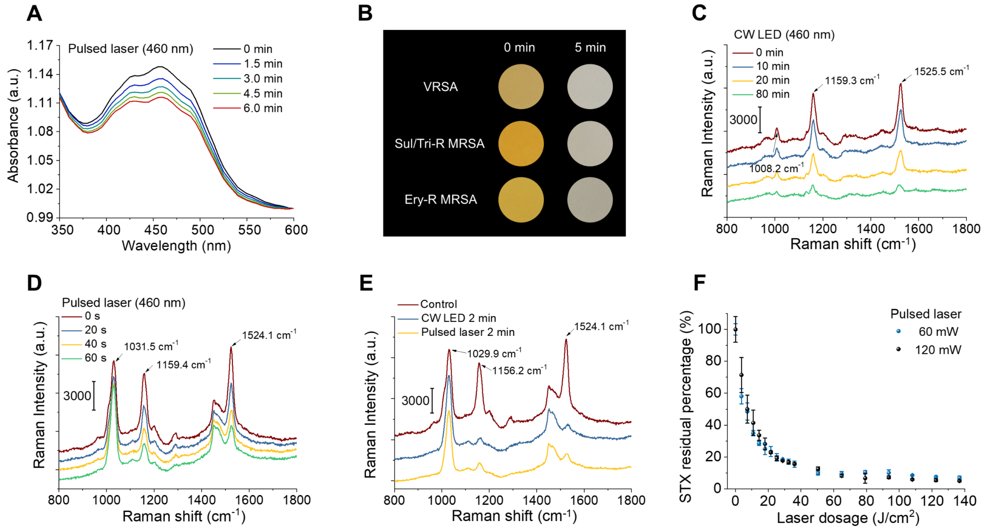


**Fig. S1. Photophysics and photochemistry of pulsed laser photolysis of STX.** (A) Absorption spectroscopy of MRSA solution over 460 nm nanosecond pulsed laser treatment time. All measurements were performed on the same sample. (B) Digital images of bacterial colonies of multidrug-resistant *S. aureus* isolates before and after 5 min laser treatment (460 nm). Bacterial strains include vancomycin-resistance *S. aureus* (VRSA), sulfamethoxazole/trimethoprim-resistant MRSA (Sul/Tri-R MRSA), and erythromycin-resistant MRSA (Ery-R MRSA). Image were recorded with sample sandwiched between two transparent glass cover slides over a black paper. (C) Resonance Raman spectroscopy of MRSA colony over 460 nm CW LED treatment time (measured on the same colony). Numbers indicate major Raman peak positions. (D) Resonance Raman spectroscopy of STX solution over 460 nm nanosecond pulsed laser treatment time (measured on the same colony). Numbers indicate major Raman peak positions. These peak positions at 1031 and 1524 $cm^{-1}$ and the peak amplitude change at 1031 $cm^{-1}$ are different from that of MRSA colony, indicating different chemical environment for STX in extract solution and MRSA membrane. (E) Resonance Raman spectroscopy of STX solution by nanosecond pulsed laser and CW LED under the same illumination power, area, and center wavelength (460 nm) highlighting a similar STX photolysis efficiency. STX solution were extracted directly from MRSA cells. (F) STX photolysis kinetics of MRSA colony by nanosecond pulsed laser under the same dosage but different illumination power (460 nm). CW, continuous wave. N=3 for all the above measurements.

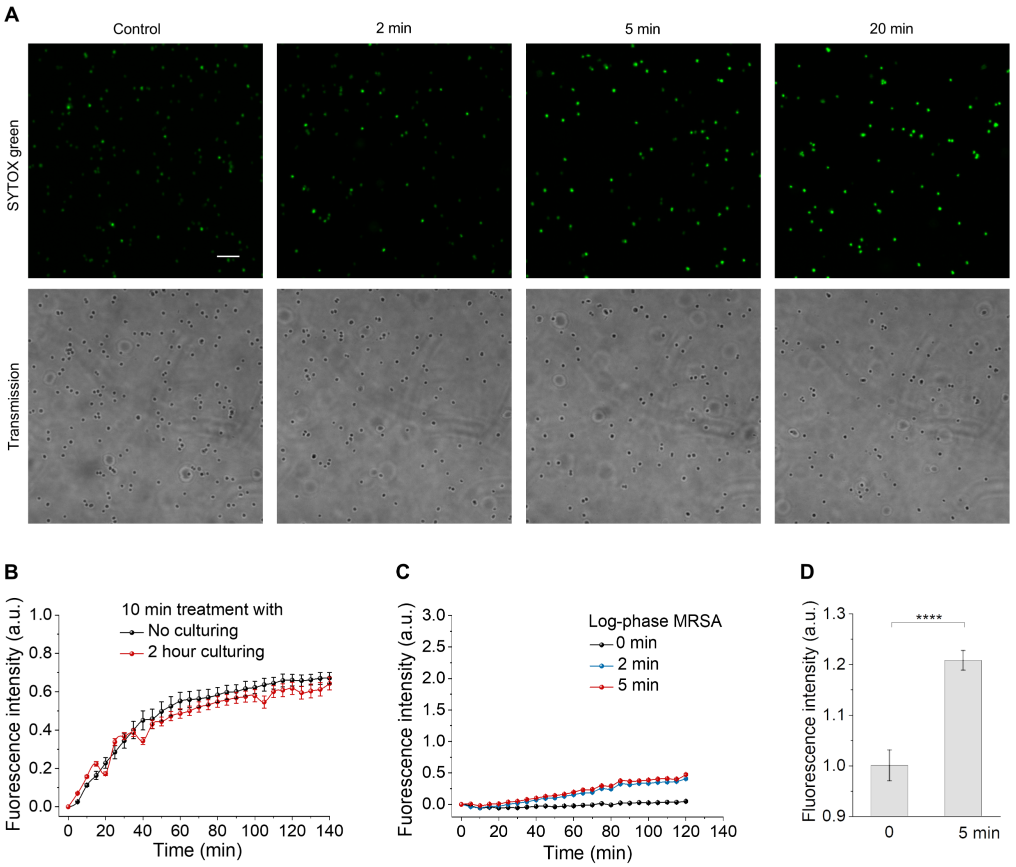


**Fig. S2. First mechanism for photo-disassembly of membrane microdomains: membrane permeabilization.** (A) Confocal fluorescence images of intracellular uptake of SYTOX green by stationary-phase MRSA cells with or without pulsed laser treatment showing STX photolysis-mediated SYTOX green uptake. (Top) fluorescence images. (Bottom) corresponding transmission images. Scale bar, 5 µm. (B) Real-time intracellular uptake kinetics of SYTOX green by stationary-phase MRSA after 10 min pulsed laser treatment with or without followed by 2-hour culturing. (C) Real-time intracellular uptake kinetics of SYTOX green by log-phase MRSA with or without pulsed laser treatment. (D) Fluorescence detection of gentamicin-Texas red uptake by stationary-phase MRSA with or without pulsed laser treatment. N=3 for all the above measurements.

**A** Daptomycin-BODIPY

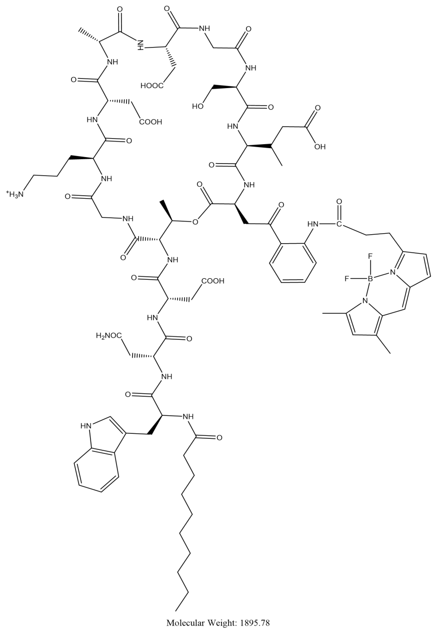


**Fig. S3. Second mechanism for photo-disassembly of membrane microdomains: membrane fluidification.** (A) Molecular structure of daptomycin-BODIPY.

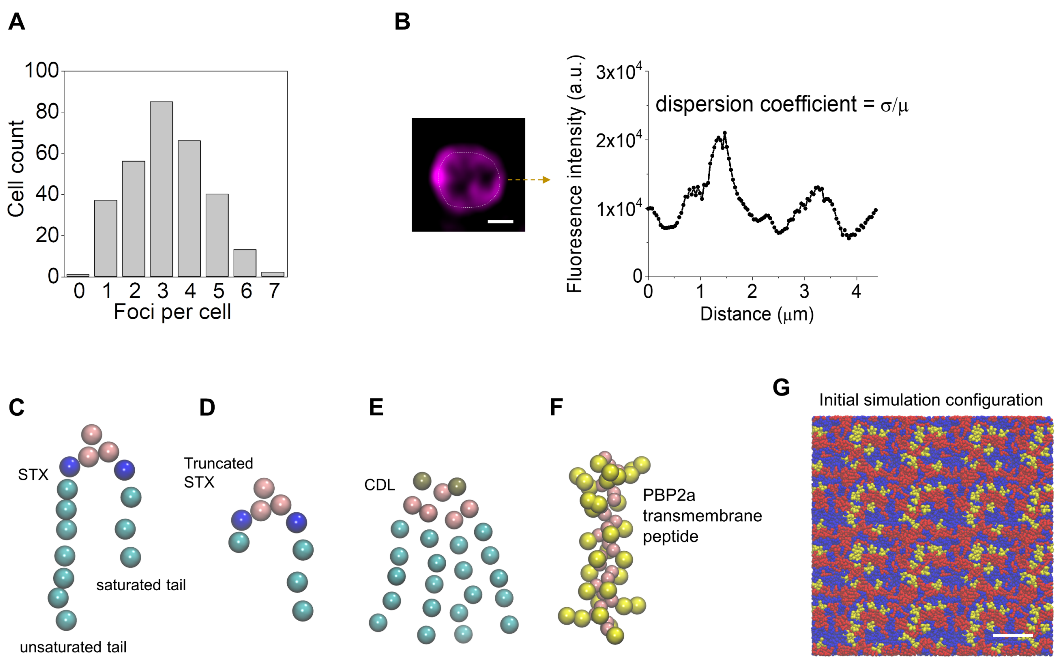


**Fig. S4. Third mechanism for photo-disassembly of membrane microdomains: membrane protein detachment.** (A) Statistical analysis of foci number on stationary-phase MRSA cells with $N \geq 300$. (B) Quantification of PBP2a dispersion by calculating coefficient of variation from each cell from standard deviation ($\sigma$) and mean ($\mu$) of the plotted signal intensity. Scale bar, 0.5 µm. (C-F) Coarse-grained representations of the (C) full-length STX, (D) truncated STX, (E) cardiolipin (CDL), and (F) PBP2a transmembrane peptide. (G) Initial configuration of modeled membrane. Full-length STX lipids, red; cardiolipin lipids, blue; PBP2a peptides, yellow. Water and ions are made invisible for clarity. Scale bar, 5 nm.

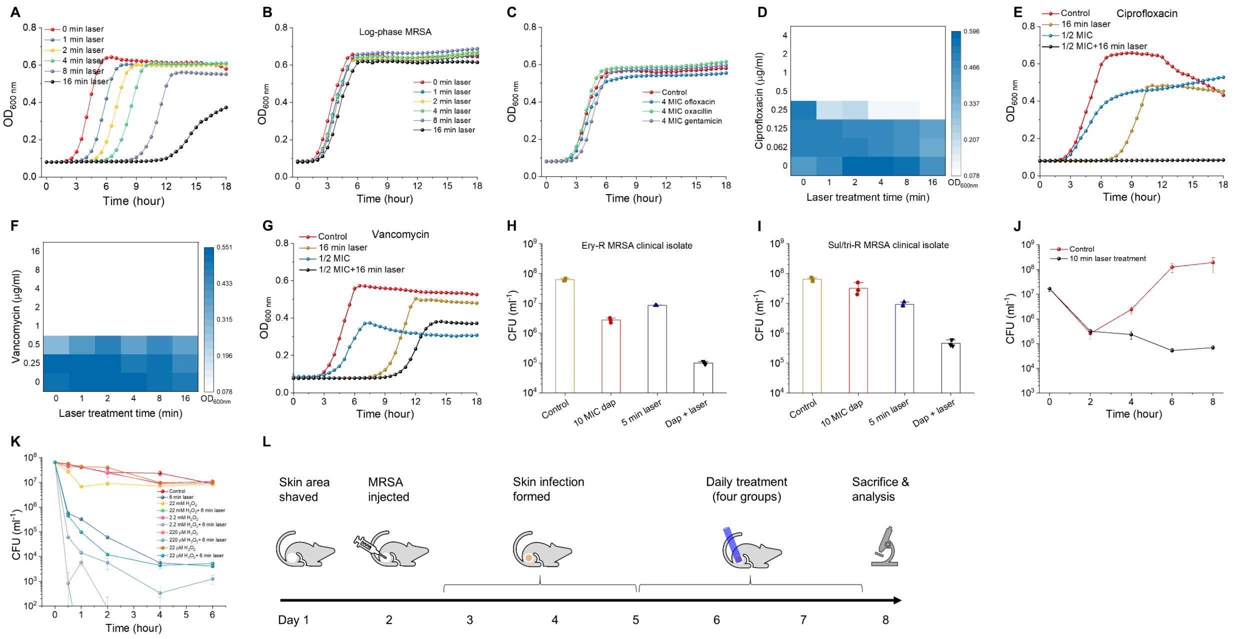


**Fig. S5. Photo-disassembly of membrane microdomains potentiates a broad spectrum of conventional antibiotics.** Post-exposure effect of (A) stationary-phase and (B) log-phase MRSA cells after different laser treatment time highlighting that the growth delay induced by laser treatment is dependent on STX abundance in MRSA cells. (C) Post-antibiotic effect of stationary-phase MRSA cells for ofloxacin, oxacillin, and gentamicin relative to the control. (D-G) Checkerboard assay results for synergy evaluation between laser treatment and different classes of antibiotics: (D, E) ciprofloxacin, (F, G) vancomycin. (E, G) Selected cell growth curves acquired from corresponding checkerboard assay results for each antibiotic. Time-dependent killing of stationary-phase (H) sulfamethoxazole/trimethoprim-resistant MRSA (Sul/Tri-R MRSA) and (I) erythromycin-resistant MRSA (Ery-R MRSA) in phosphate-buffered saline for four different treatment groups. (J) Time-dependent killing of stationary-phase MRSA in fresh human whole blood with or without 10 min laser treatment. (K) Time-dependent killing of stationary-phase MRSA in phosphate-buffered saline supplemented with different concentration of hydrogen peroxide after different laser treatment time. (L) Schematic of experiment design for mice skin infection model. N=3 for checkerboard assay of each antibiotic. N=3 for CFU enumeration.

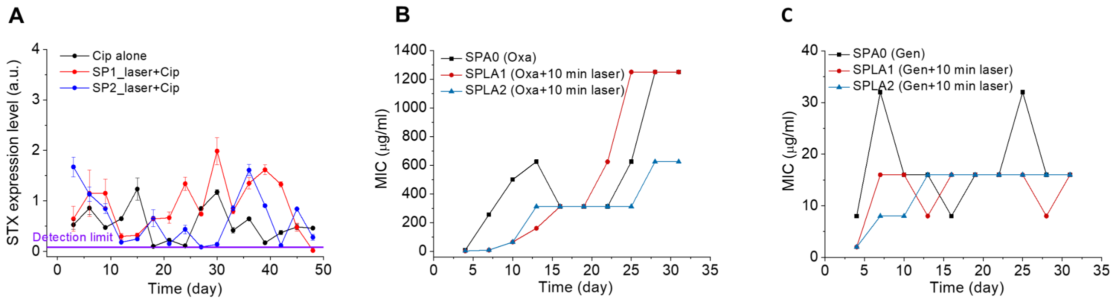


**Fig. S6. Photo-disassembly of membrane microdomains inhibits resistance development to conventional antibiotics.** (A) STX abundance in stationary-phase MRSA cells over 48-day serial passage in the presence of sub-MIC levels of ciprofloxacin with or without 10 min laser treatment. quantified via Raman peak amplitude at 1161 $cm^{-1}$. Resistance acquisition over 48-day serial passage in the presence of sub-MIC levels of antibiotics with or without 10 min laser treatment: (B) oxacillin, (C) gentamicin.

| Clinical isolates | Antibiotics | MIC (µg/ml) | Source/description |
|---|---|---|---|
| VRSA NR-46419 (VRSA 9) | Vancomycin | 256 | Isolated in 2007 in Michigan, USA from a left plantar foot wound of a 54-year-old female, who recently received a 4-week course of vancomycin and levofloxacin to treat osteomyelitis of the left metatarsals. |
| MRSA NRS384 (MRSA USA 300) | Erythromycin | 64 | Isolated from a wound in Mississippi, USA. It is a community-acquired MRSA strain. |
| MRSA NRS385 (MRSA USA 500) | Sulfamethoxazole/trimethoprim | 256 | Isolated from a bloodstream sample in Connecticut, USA. It is a hospital-acquired MRSA strain. |

| Bacterial strain | Antibiotics | MIC (µg/ml) |
|---|---|---|
| MRSA USA 300 | Daptomycin | 8 |
| | Gentamicin | 8 |
| | Oxacillin | 8 |
| | Ofloxacin | 0.5 |

**Table S1.** Minimum inhibitory concentrations of selected antibiotics against the tested bacterial strains. N=3 for each measurement.
